# Identification of circRNAs linked to Alzheimer’s disease and related dementias

**DOI:** 10.1101/2022.07.13.499937

**Authors:** Sambhavi Puri, Junming Hu, Zhuorui Sun, Mintao Lin, Thor Stein, Lindsay Farrer, Benjamin Wolozin, Xiaoling Zhang

## Abstract

We conducted a circular-transcriptome-wide analysis and examined circRNA expression patterns associated with AD and clinical and neuropathological AD severity measures in human hippocampus and cortex brain regions. We validated prior studies of circRNA in AD cortex, and demonstrate novel patterns of expression in the AD hippocampus. We also examined circRNA expression across multiple types of dementia and show that circRNA expression differs by dementia subtype. We demonstrate robust circRNA expression in human neuronal precursor cells (NPCs). Then, using NPCs to study qPCR validated circRNA, we show that exposure to oligomeric tau elicits downregulation of circRNA similar to that observed in AD brain. These data identify circRNA that are changed with AD, and suggest that treating neuronal cells with oligomeric tau can recapitulate some of these changes.

## Background

Late-onset Alzheimer’s disease (AD) is a complex neurodegenerative disorder that is the most common cause of dementia. Recent genomic studies have discovered thousands of well-expressed, stable circular RNAs (circRNAs)^1^. These circRNAs are produced from both protein-coding genes and non-coding regions of the genome via a process known as back-splicing^2-4^. CircRNAs are more enriched in neuronal tissues and are often derived from genes specific for neuronal and synaptic function^3,5-7^. The discovery of circRNAs opens an entirely new window into mechanisms of neurodegeneration in Alzheimer’s disease (AD) and related dementia (ADRD).

CircRNAs could contribute to neurodegeneration by acting as sponges that sequester microRNA and RNA-binding proteins (RBPs) away from normal mRNA targets, altering the splicing or expression of target mRNAs^8^. RBPs also regulate circRNA production by binding to the flanking intronic sequences of circRNAs, which contain many conserved binding sites of splicing factors/RBPs^9-11^. Thus, sequestration of RBPs in protein aggregates could cause dysfunctional regulation of circRNAs. Studies from the field of oncology reveal wide ranging effects of circRNAs on tumor biology^12^. Biomarker studies of circRNA also demonstrate multiple examples of associations with tumor levels, type, prognosis or therapeutic response.

The expression pattern of circRNA in AD is only beginning to be studied^1^. Studies have begun to characterize circRNA expression in AD brain and mouse models of AD ^13,14^. However, numerous critical questions remain. Methods for detecting circRNA are evolving, the effects of disease heterogeneity on circRNA expression are poorly understood, and comparisons between brain regions and the regulation of circRNA by AD-related factors in human neurons are unexplored. Thus, the field of circRNA presents a major unexplored avenue of RNA metabolism that demands investigation to expand our understanding of AD mechanisms.

In this study, we conducted a circular-transcriptome-wide analysis and examined circRNA expression patterns associated with AD and clinical and neuropathological AD severity measures in human hippocampus and cortex brain regions. We validate prior studies of circRNA in the AD cortex, and demonstrate novel patterns of expression in the AD hippocampus. We also show that circRNA expression differs by dementia subtype. We demonstrate robust circRNA expression in human neuronal precursor cells (NPCs). Using NPCs to study qPCR validated circRNA, we show that exposure to oligomeric tau elicits downregulation of circRNA similar to that observed in AD brain.

## Results

### Study design/Subject Characteristics

The study design included calling and quantifying circRNA in two brain regions from three study cohorts in which independent RNA-seq datasets were derived from neuropathologically confirmed AD case and control brain tissues. In the hippocampus dataset, 210 individuals were assessed at the ADRC of Boston University School of Medicine, and their demographic, clinical severity and neuropathological information are presented in Table 1. Out of 210 BU ADRC participants, 207 European ancestry individuals remained for inclusion in the analysis. These include 79 Pure AD cases, 56 AD cases with Vascular Dementia and 47 AD cases with Lewy body dementia and 21 cognitively normal controls. Table 1 presents the clinical characteristics of the study participants including CDR, Braak score as well as CERAD score (P<0.05). Table 1 also presents subject age (mean=81.0, P=9.15), sex distribution (P=0.00037) and RNA integrity number (RIN).

**Table 1.** **Distribution of subjects in BU-ADRC Hippocampus data including** clinical and neuropathological measurement

| Distribution of subjects in BU Hippocampus data (n=207) |  |  |  |  |  |
| --- | --- | --- | --- | --- | --- |
| Variable | AD_pure | AD+VaD | AD+LBD | AD other | Control |
| **N** | 79 | 56 | 47 | 4 | 21 |
| **Age(yr)**<br>**(SD)** | 76.9<br>(8.30) | 84.6<br>(9.70) | 79.9<br>(5.85) | 80.3<br>(5.56) | 89.0<br>(9.00) |
| **Female**<br>**N (%)** | 12<br>(15.19) | 17<br>(30.36) | 7<br>(14.89) | 0<br>(0) | 10<br>(47.62) |
| **RIN** |  |  |  |  |  |
| **≤4.3** | 24 | 12 | 12 | 2 | 4 |
| **4.3<x≤5.6** | 18 | 19 | 14 | 1 | 1 |
| **5.6<x≤6.9** | 16 | 14 | 14 | 1 | 5 |
| **>6.9** | 21 | 11 | 7 | 0 | 11 |
| **CDR** |  |  |  |  |  |
| **0** | 0 | 0 | 1 | 9 | 0 |
| **0.5** | 1 | 0 | 2 | 3 | 0 |
| **1** | 1 | 1 | 5 | 2 | 0 |
| **2** | 1 | 3 | 11 | 0 | 0 |
| **3** | 30 | 16 | 30 | 3 | 0 |
| **Braak** |  |  |  |  |  |
| **1** | 0 | 0 | 0 | 0 | 4 |
| **2** | 0 | 0 | 0 | 0 | 11 |
| **3** | 1 | 3 | 2 | 0 | 5 |
| **4** | 3 | 5 | 5 | 3 | 0 |
| **5** | 26 | 21 | 17 | 1 | 1 |
| **6** | 49 | 27 | 23 | 0 | 0 |
| **CERAD** |  |  |  |  |  |
| **0** | 0 | 0 | 0 | 0 | 0 |
| **1** | 70 | 46 | 39 | 3 | 2 |
| **2** | 7 | 7 | 8 | 1 | 2 |
| **3** | 2 | 3 | 0 | 0 | 8 |
| **4** | 0 | 0 | 0 | 0 | 9 |

N:sample size; RIN: RNA integrity number; SD: standard deviation. Values for Age and RIN expressed as mean ± sd.

For comparison to a different brain region, an independent, publicly available, Accelerating Medicines Partnership–Alzheimer’s Disease (AMP-AD) rRNA-depleted RNA seq data for inferior frontal gyrus tissue (BM44) generated by Mount Sinai Brain Bank (MSBB) (216 AD cases, 62 controls) was analyzed using the same pipeline (Supplementary Table 1). Additional hippocampus RNA-seq data of European ancestry generated by the Adult Changes in Thought (ACT) were downloaded from NIAGADS (25 AD cases and 50 Control) and analyzed using the same workflow (Supplementary Table 2).

### CircRNA profiling/expression in hippocampus and cortex of human aging/AD brains selects for circRNA from synaptic cognate genes

rRNA-depleted RNA from 210 hippocampi of subjects from ADRC were sequenced and evaluated for quality control. Samples with poor quality RNA or outliers were excluded, leaving 207 samples for inclusion in the analysis. The resulting hippocampal sequencing data yielded an average 45 million paired-end reads per individual. Reads were aligned to the human reference genome sequence (GRCh38) using STAR (version 2.6.1c) and CIRCexplorer 3 was used to identify back-splice junctions and call circRNAs. As shown in Supplementary Table 3, in the hippocampus (BU ADRC dataset), 4,092 circRNAs from 2,056 unique gene locus were detectable (FPB>0 in >50% of the samples, FPB=fragments per billion mapped base). In the cortex BM44 region (MSBB AMP-AD dataset), 4,912 circRNAs were detectable from 2,414 unique gene locus with an average of 2 circRNA isoforms per gene.

Next, we selected for circRNAs that were widely expressed by selecting those circRNAs that exhibited expression in over 75% samples at FPB>0. More stringent expression thresholds progressively reduced the number of circRNA reaching significance. Application of this expression threshold reduced the number of circRNAs by more than 50% (1,762 and 2,212 in the hippocampus and cortex, respectively). At FPB>1, 285 hippocampal circRNAs and 980 cortical circRNAs were expressed in over 75% samples, and at FPB> 2 only 114 circRNAs in the hippocampus vs. 512 in the cortex. The smaller fraction of highly expressed circRNAs observed in the hippocampus versus cortex might reflect the lower RNA quality in the hippocampal samples compared to the cortical samples. These findings are consistent with prior observations that the expression levels of non-coding RNAs including circRNAs are much lower than those of protein-coding mRNAs. All circRNAs expressed at >= 1 FPB in over 75% samples of hippocampus and cortex are listed in Supplementary Tables 4 and 5 in the online-only data supplement, respectively.

Gene ontology analysis of hippocampal circRNA showed that categories of synaptic and pre-synaptic localization, and synaptic transmission were enriched in the 232 circRNA host genes expressed moderately (FPB>1 in over 75% samples) and 103 expressed highly (FPB>2) in hippocampus (Supplementary Figure 1). In addition, we compared circRNAs detected from the hippocampus to MSBB BM44 cortex data. We found the percentage of commonly expressed circRNAs in both brain regions is decreased, while the percentage of unique circRNAs in cortex region is increased (Supplementary Table 3).

### Differential expression analysis to identify circRNA signatures for AD in hippocampus and cortex

As shown above, a comparable number of circRNAs were detected at FPB>0 in the hippocampus (n=4,092) and in the cortex (n=4,912) with 80.5% (3,295/4,092) expressed in over 50% of both brain region samples. Therefore, for the differential expression (DE) analysis, we included circRNAs detected (FPB>0) across 50% of our BU ADRC hippocampus samples and MSBB BM44 cortex samples. Although more than one circRNAs was detected from each gene locus, for this study we selected the most abundant/significant circRNA from each gene locus. We obtained a total of 2,258 circRNAs, then meta-analyzed with the same circRNA transcripts in the MSBB-BM44 cortex dataset. At a false discovery rate (FDR) <0.05, we found 48 circRNAs (one circRNA per gene) differentially expressed between AD and controls (Table 2), with 9 circRNAs expressed only in the cortex. Among these, the expression patterns for 38 circRNAs were in the same direction for the hippocampus and BM44 cortex regions with 3 up-regulated and 35 down-regulated. Notably, circQKI was highly expressed (average FPB=2.3) and up-regulated in both AD brain regions (logFC=0.22, p=2.23 ×10^−2^ in cortex, and logFC=0.61, p=6.79 ×10^−3^ in hippocampus). circQKI is a RBP, reported to bind to the intron loop to regulate circRNA biogenesis in epithelial-mesenchymal transition tissue/cells^10^ (Figure 2A-B). circMAN2A1 was overexpressed in both AD brain regions although only significantly different in the cortex (Table 2), which may due to a small number of control samples in our hippocampus dataset. No circRNAs exhibiting significant differential expression between AD and control showed opposite disease-linked changes between the cortex and hippocampus.

**Table 2.**
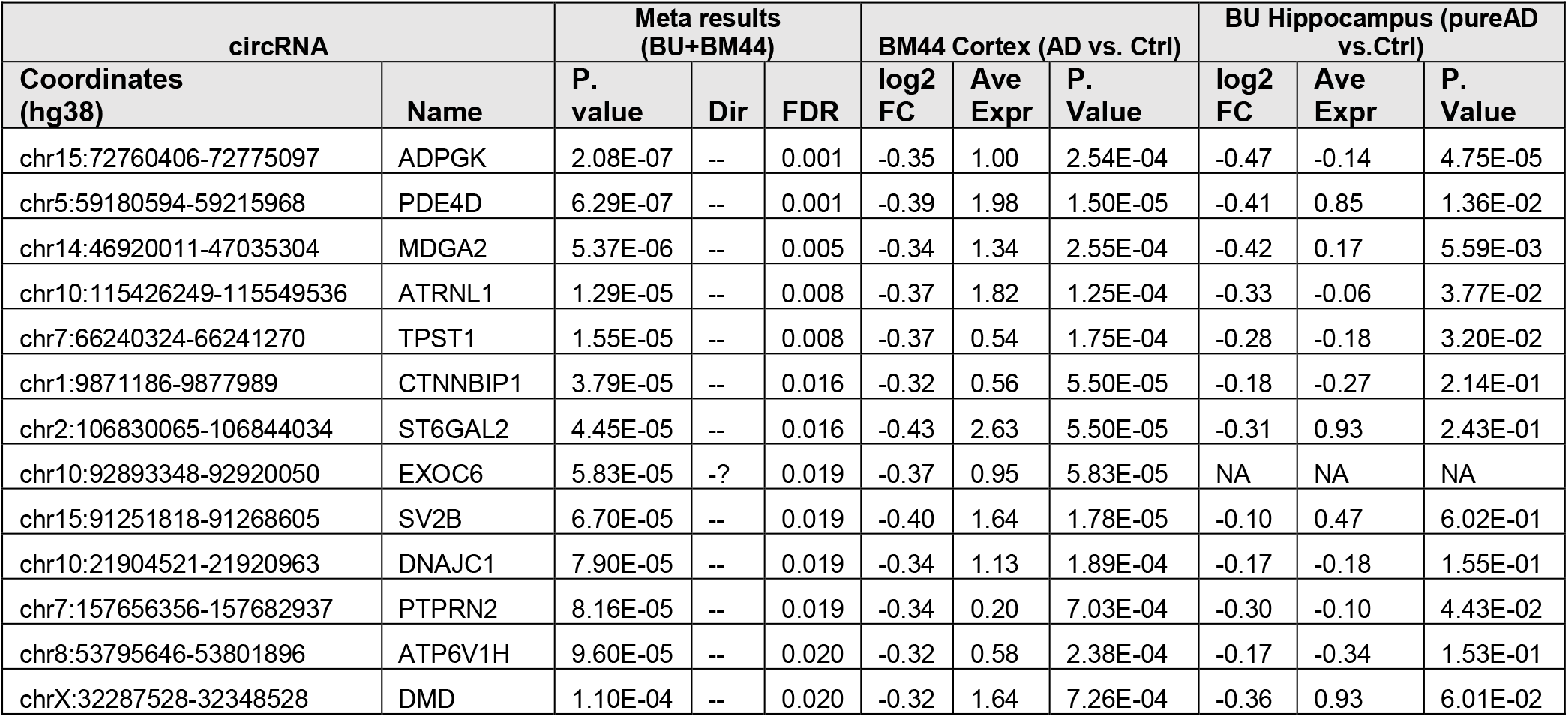
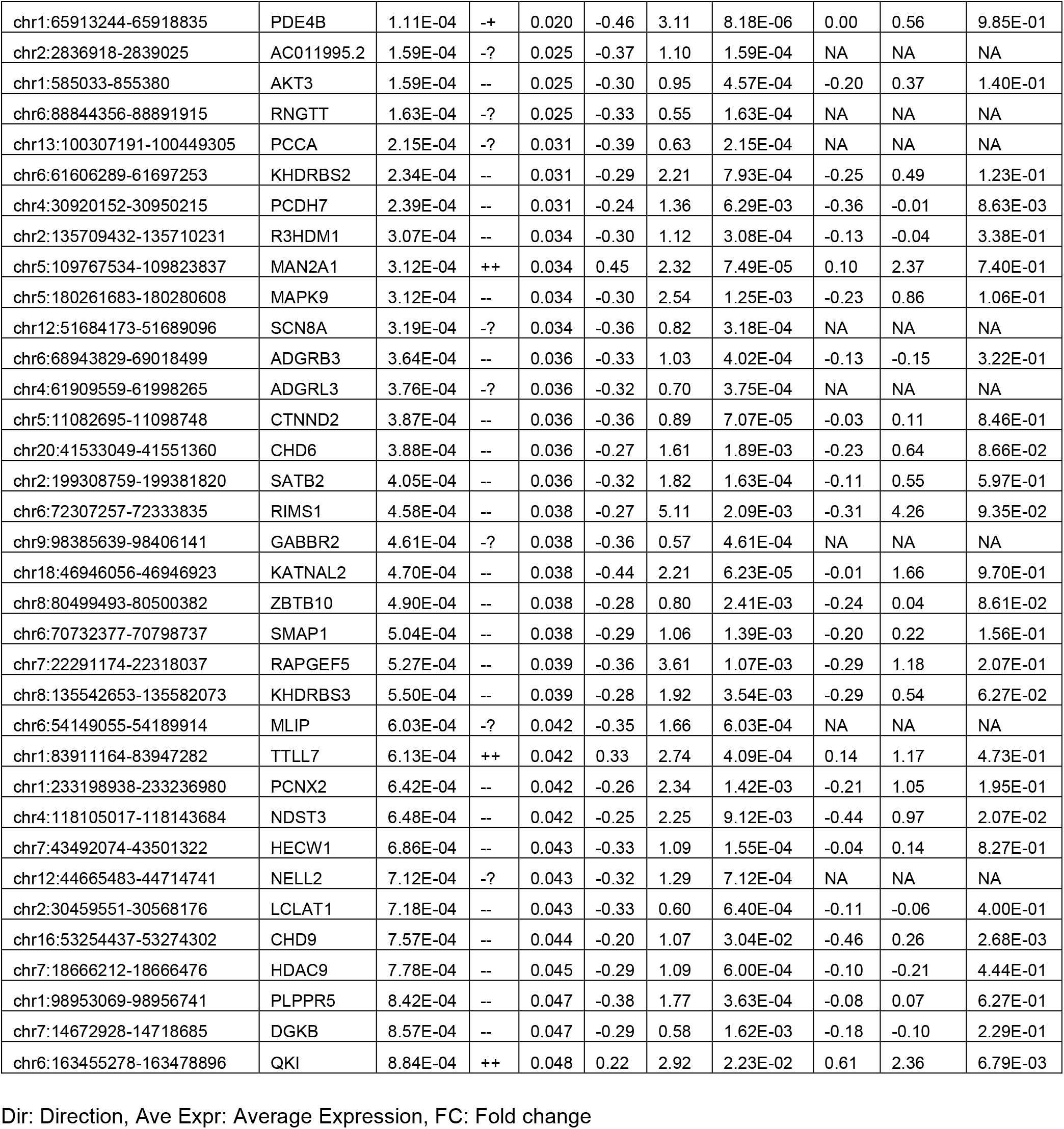
Meta-analysis of circRNA differentially expressed between AD and controls in the hippocampus and cortex at FDR<0.05 (n=48).

**Figure 1.**
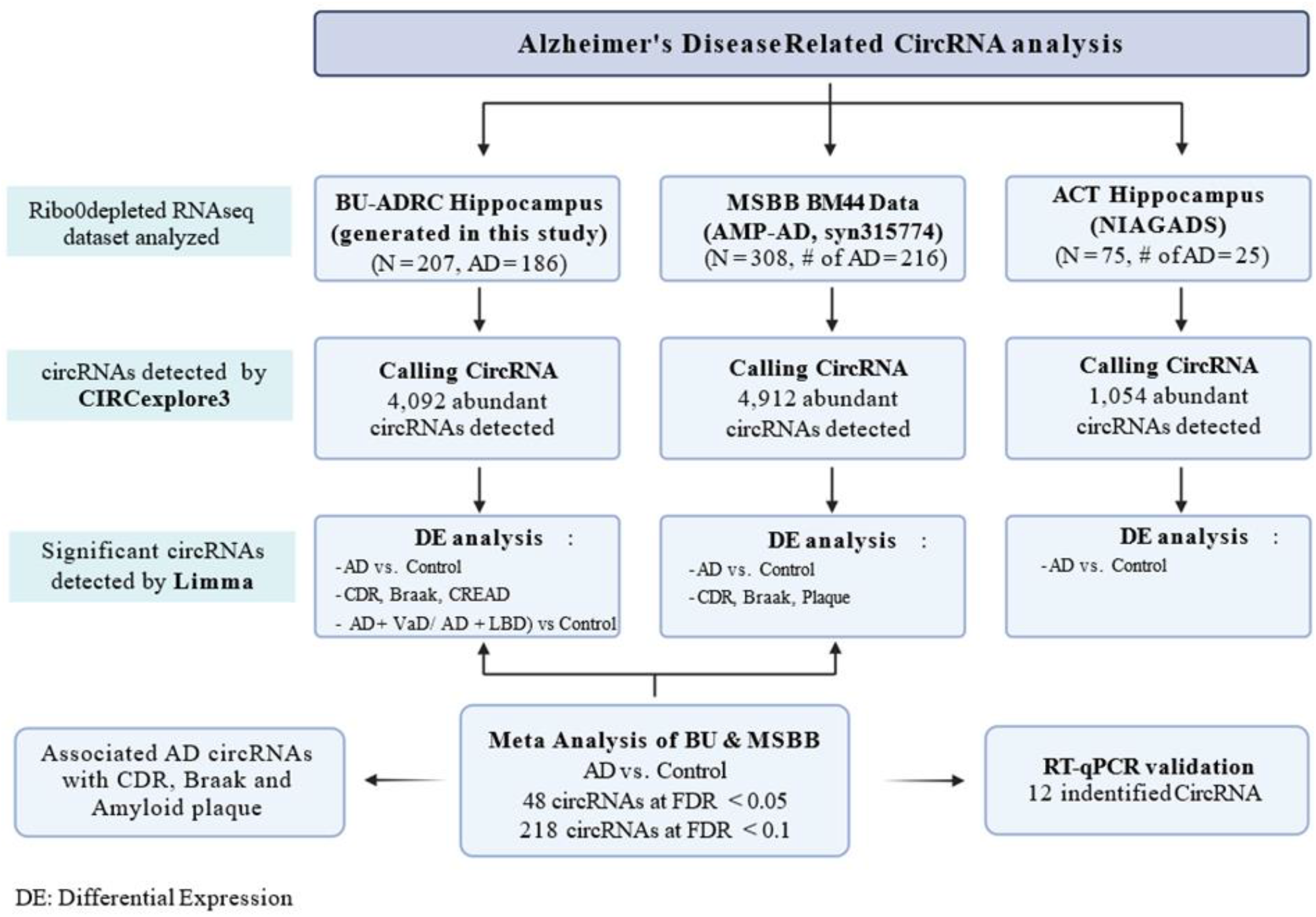
Flowchart represents the pipeline for circRNA detection and differential expression analysis in three AD brain RNA-seq data.

**Figure 2.**
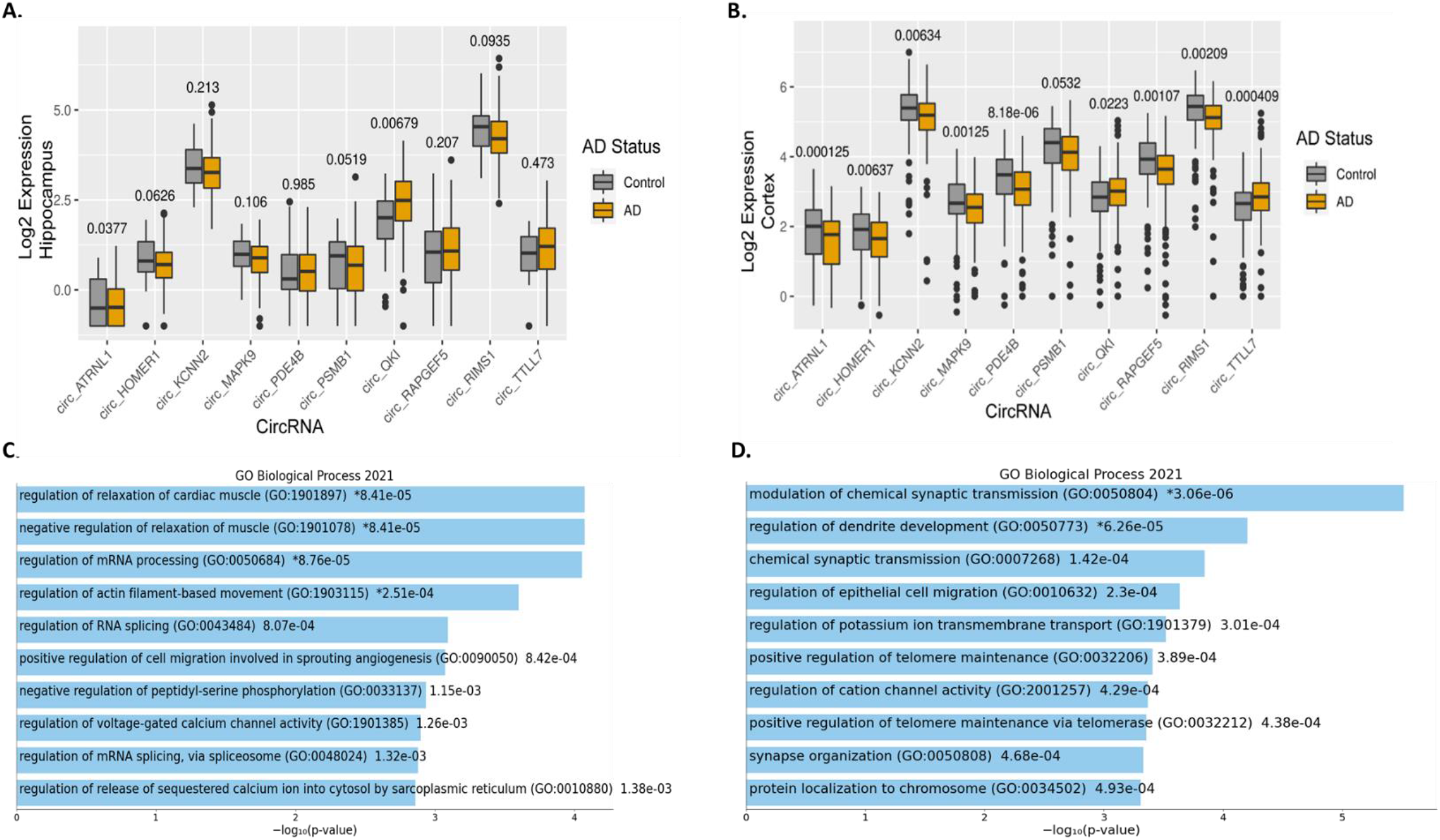
Top differentially regulated circRNAs between AD and control. (2A-B) Boxplots showing the expression levels (represented as log2 transformed FPB) and distribution of top differentially regulated circRNA in the hippocampus (BU-ADRC) and BM44 cortex (MSSB AMP-AD), respectively. (**2C-D)** GO pathways enrichment analysis of cognate mRNA genes for 48 and 218 circRNAs, respectively that are differentially expressed at FDR<0.05 and FDR<0.1, respectively.

We proceeded to check these 48 circRNAs in an independent hippocampus dataset (ACT). Among were 18 circRNAs detected (FPB>0) across 50% of the ACT samples. Three circRNAs (MAPK9, RAPGEF5, NDST3) were also significantly down-regulated in AD. circQKI was again expressed relatively highly (FPB=2.14) and up-regulated (logFC=0.44, p=0.015) in the AD hippocampus (see Supplementary Table 6 for 218 circRNAs at FDR<0.1 with results from all 3 datasets analyzed for AD status). Of note, circHOMER1, a neuronal-enriched circRNA abundantly expressed in the brain, was more abundant in the cortex compared to the hippocampus, and down-regulated in AD across all three datasets with a similar fold change (logFC= -0.27 in cortex, logFC= -0.28 in two hippocampus datasets) (Figure 2A-B).

We proceeded to use the corresponding host/cognate mRNA genes for the 218 differentially expressed circRNAs (FDR<0.1) to perform GO/KEGG enrichment analysis (Figure 2C, Supplementary Table 7). The cognate mRNA transcripts showed significant enrichment for the synapse, postsynaptic density (17/209 genes, FDR = 4.07e-06), synaptic vesicle membrane (8/209 genes, FDR=2.34e-03), axon and neuronal cell body (13/209 genes, FDR=2.71e-02) pathways. Further, enrichment for RNA processes and RNA splicing were observed among the 48 differentially regulated circRNAs (3/36 genes, FDR=8.76 e-05).

### Top differentially expressed circRNAs correlate with AD pathology and cognitive loss/CDR in the hippocampus and cortex

Next, we sought to understand the relative importance of clinical and pathological outcomes in the association with circRNA expression patterns. circRNA expression was correlated with clinical dementia rating (CDR) and neuropathological AD-related traits including CERAD neuritic plaque score and Braak stage in both the hippocampus and cortex regions. CDR measures the severity of cognitive impairment ranging from no dementia (score=0) to severe dementia (score=1). Braak stages in AD are defined by the distribution of neurofibrillary tangles in the brain. Stages I and II involve confinement of neurofibrillary tangles to the transentorhinal region, the tangles spread to limbic regions in stages III and IV and the neocortex in stages V and VII. Neuritic plaque comprises amyloid-beta deposits and deteriorating neuronal material surrounding amyloid-beta deposits. CERAD neuritic plaque score ranges from 1 (definite AD) to 4 (no AD). As shown in Supplementary Table 8. Most circRNAs were significantly associated with AD case status, dementia severity and neuropathological severity common to both hippocampus and cortex, while some circRNA showed phenotypic correlations that were selective for the cortex or hippocampus.

For 48 circRNAs differentially expressed between AD and normal control brains (FDR<0.05), we checked their association with Aβ pathology, tau pathology and clinical symptom of dementia scores. As shown in Figure 3A, among 45 circRNAs with association results for CDR, CERAD and Braak score, 69% (31/45) circRNAs were significantly associated with all three traits, 37/45 correlated with cognition (CDR), 42/45 correlated with Braak score, and 39/45 correlated with plaque Aβ-load (CERAD). More AD-associated circRNAs correlated between CDR and Braak than CDR and Plaque (80% vs. 71%), indicating the critique role of Braak/tau-load with clinical diagnosed dementia stage and neuropathologically confirmed dementia stage. In addition, 6.7% (3/45) circRNAs exhibited an expression pattern correlating with Aβ pathology, while 13.3% (6/45) circRNAs exhibited an expression pattern correlating with tau pathology. Similar patterns were observed for 218 circRNAs differentially expressed between AD and control (Supplementary Figure 3). For selected circRNAs (circRIMS1, KCNN2, PSMB1, ATRNL1, PDE4B, HOMER1), scatterplots of their expression in hippocampus and cortex vs. CDR/Plaque/Braak are shown in Figure 3B. Each of these circRNA show significant correlation with Braak and CDR scores in the cortex. The cognate linear transcripts are associated with synaptic functions for *RIMS1, KCNN2* and *HOMER1* ^15,16^.

**Figure 3.**
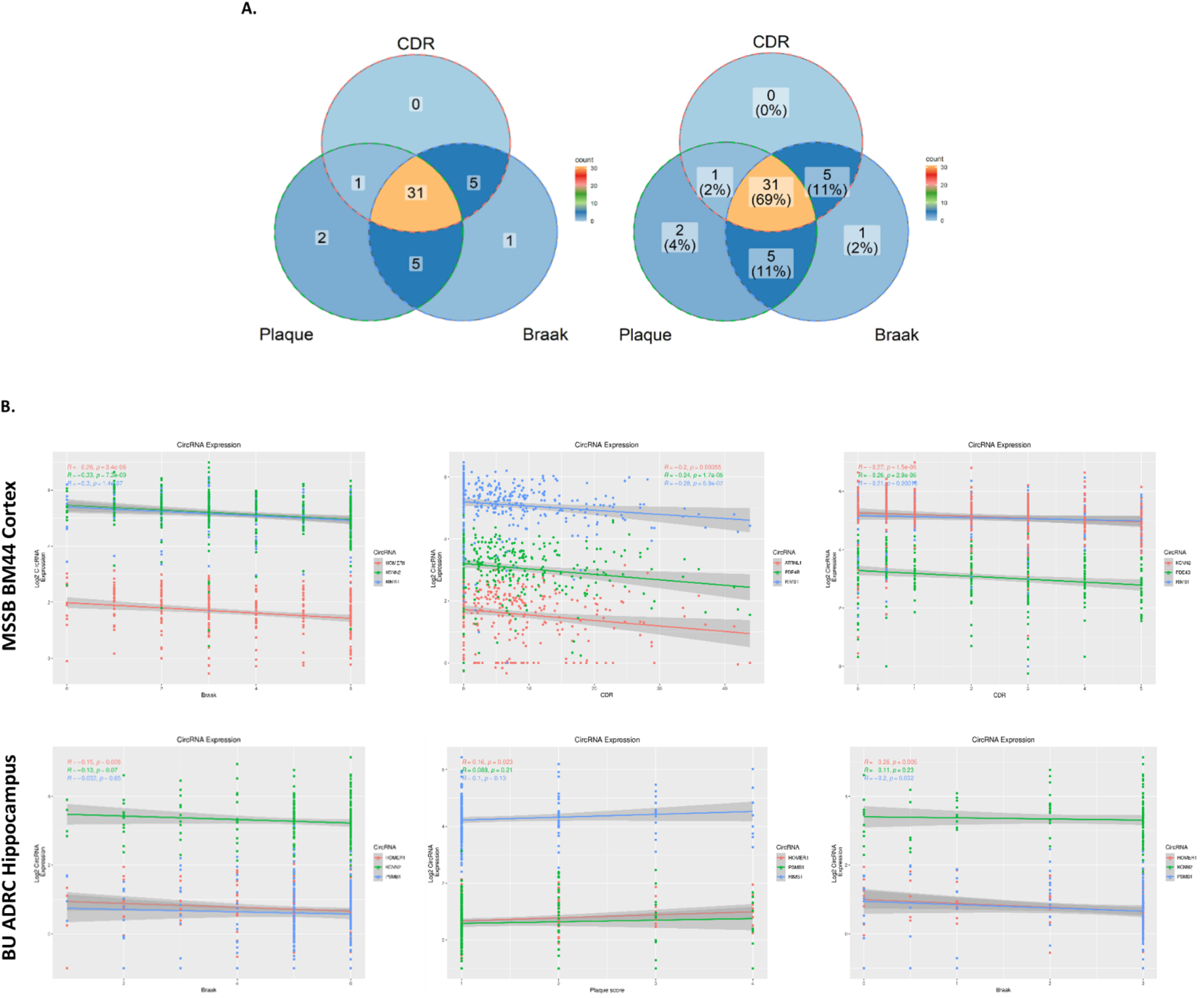
CircRNA associated with different indices of AD. (A) Venn Diagram depicting association of 48 DE circRNAs with AD pathology (Braak and Plaque) and CDR. (B) Scatter plot showing the association of CDR, Plaque score, and Braak score with the top three significant circRNA. Coordinates are shown as color-coded dots for different kinds of circRNA and their correlation across subjects (within AD and control) is represented by fitted solid lines. Note, plaque mean score in MSBB BM44 is positively correlated with amyloid severity, but it is vice-versa in BU-ADRC hippocampus which is a CERAD score.

### The pattern of circRNA differential expression in AD cases is modified by the presence of vascular or Lewy body pathology (AD+VaD, AD+LBD)

Patterns of circRNA expression were examined from a range of dementia cases, including AD, Lewy body disease (LBD) and vascular dementia (VaD). Cases of AD frequently show multiple types of pathology, such as vascular pathology and Lewy body pathology. To explore the impact of other pathologies on circRNAs we identified circRNAs expressed differently between normal controls and AD patients exhibiting pathology for vascular dementia (AD+VaD) or Lewy body dementia (AD+LBD). As shown in Figure 4A, at P<0.05, only 39 circRNA exhibited differential expression that was common to all three disease conditions compared to controls, which was the smallest group of circRNAs studied. Far more circRNA exhibited expression patterns that were sensitive to pathological subtype. 82 circRNAs showed differential expression in cases of pure AD or AD, while 62 circRNAs showed differential expression patterns that were unique in AD+VaD. Interestingly, the AD+LBD group exhibited the largest number of unique differentially expressed circRNAs (n=154), even though the number of AD+LBD patients (n=47) was smaller than pure AD (n=79) and AD+VaD (n=56). This larger number of differential circRNAs associated with AD+LBD could reflect more severe neuropathological features or a stronger association with synaptic pathology. Synaptic pathology is a strong pathway for circRNAs and also a strong pathway for α-synuclein, which is the predominant aggregated protein in Lewy bodies. Examining correlations with CDR instead of disease presence (AD only) did not increase the number of circRNAs identified that exhibited differential expression patterns common among disease subvariants. Although more circRNAs are associated with CDR (n=284) compared to disease presence (AD only, n=194), only 20 circRNAs were found common among AD+VaD, AD+LBD and CDR (Figure 4B). Comparison to plaque or tangle pathology yielded similarly small amounts of overlap. Correlations with plaque pathology yielded only 14 circRNAs that were common among AD+VaD, AD+LBD (Figure 4C), while correlations with neurofibrillary tangle pathology (Braak score) yielded only 30 circRNAs that were common among AD+VaD, AD+LBD (Figure 4D). Interestingly, the largest circRNA group observed (716 circRNAs) were circRNAs exhibiting differential expression correlating with plaque pathology (Figure 4C). These results suggest that the expression of circRNA varies according to the type of neuropathology, perhaps reflecting differences in the underlying molecular regulatory pathways.

**Figure 4.**
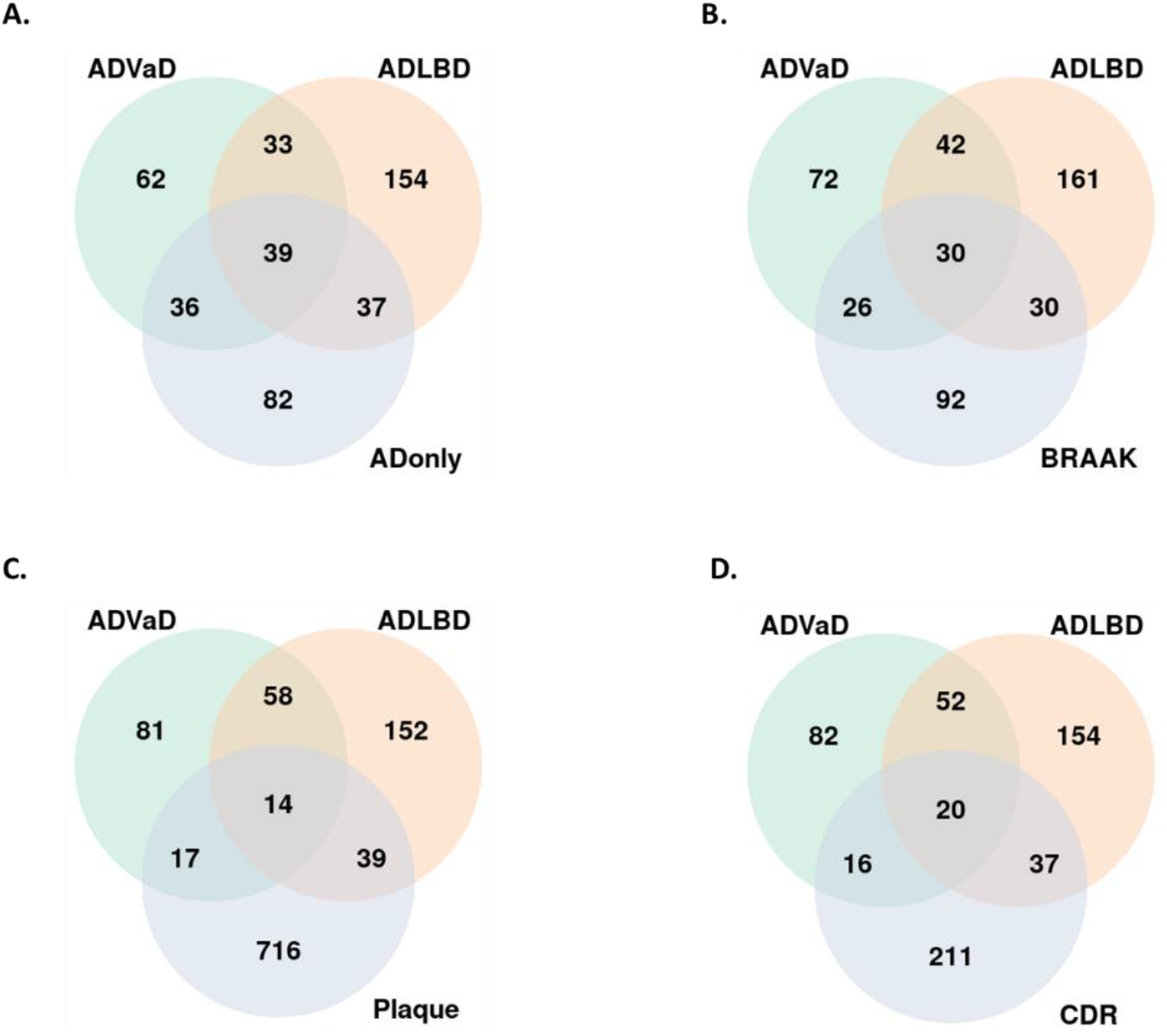
Venn Diagram of circRNAs differentially expressed in hippocampus of pure AD and AD with vascular/lewy-body neuropathy compared to elderly control brains.

### Analysis of circRNA for cognate AD-related GWAS genes or loci

To explore circRNA related to pathways most closely linked to AD, we examined circRNA for cognate genes with reported AD-related GWAS gene loci ^17^. As shown in Figure 5, circPICALM is overexpressed in the hippocampal region of AD brains (P=0.033, logFC = 0.31). Analysis of cortical BM44 tissues revealed multiple circRNAs that were differentially expressed between AD and control cases for cognate genetically linked AD gene loci, including circPICALM (P-value = 0.019, logFC = 0.21), circPSEN1(P = 0.025, logFC = 0.20), which were both upregulated in AD, and circAPP (P=0.0152, logFC = -0.21), which was down-regulated in AD. Of note, the corresponding cognate linear mRNA levels were not altered by AD status, demonstrating that the AD-associated circRNA changes in expression were independent from AD-associated changes in cognate mRNA levels.

**Figure 5.**
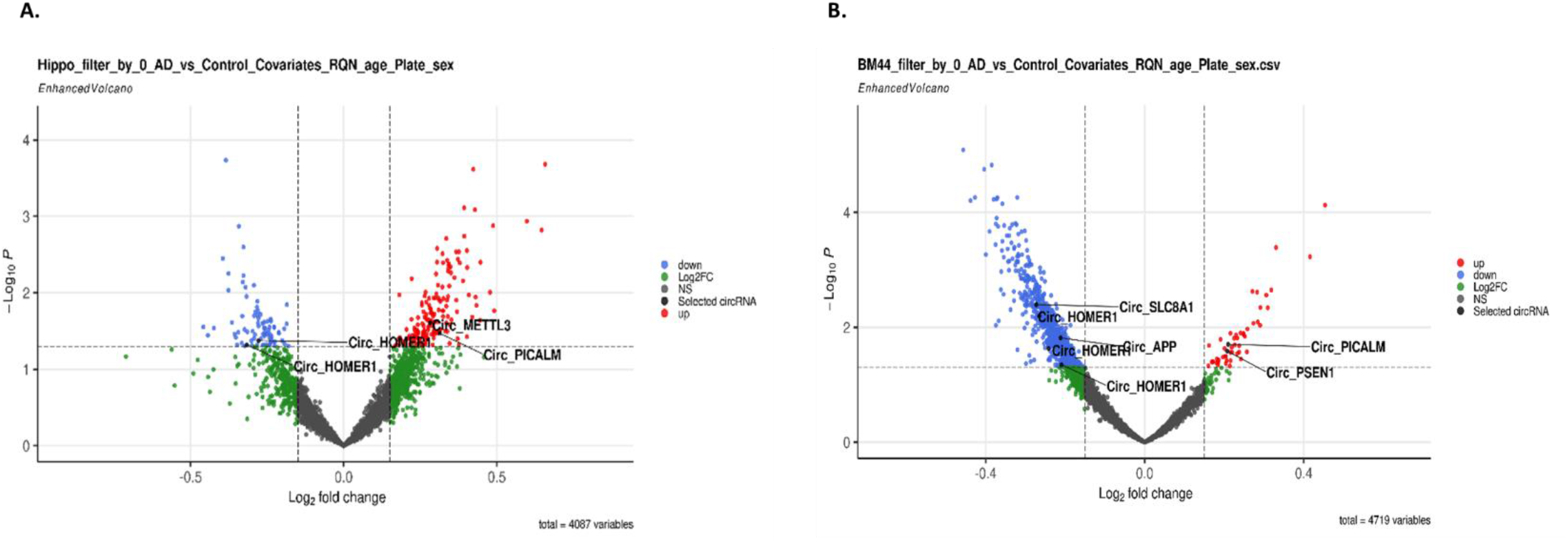
CircRNAs for known AD GWAS loci (*APP, PSEN1, PICALM*). (A-B) Volcano plots of circRNAs differentially expressed in AD vs Control in Hippocampus and Cortex brain region, respectively. P values were generated by empirical Bayes moderated test-test in limma. CircRNAs with a log-fold change > 0.15 and a −log10 adjusted P value of ≥1.3 (indicated with dashed lines) were considered significantly (red dots). Highlighted are the circRNA derived from AD-associated genes.

### RT-qPCR validation

RT-qPCR was done to validate differentially regulated circRNA in the cortex and hippocampus regions. The samples used for validation were from an independent cohort (Cortex: N=20 AD, N=20 Ctl; Hippocampus: N=18 AD, N=8, Ctrl; details in the Supplement information section). The qPCR data studies used two internal controls; GAPDH and circN4BP2L2. circN4BP2L2 (hsa_circ_0000471) was chosen specifically because it is a circRNA and has also been previously reported as a stable internal control for 22 human cell lines^18^; both informatics and qPCR analyses of our datasets (logFC=0.14, p=0.20 in hippocampus; logFC=-0.003, P=0.99 in cortex) confirmed the stability of circN4BP2L2 among the AD and control cases.

The 12 circRNAs were selected for validation based on high levels of expression in the cortex or hippocampus; analysis of circRNA with high expression levels decreased the number of PCR cycles required for detection, which therefore reduced the associated variance. The circRNAs amplified were confirmed using Sanger sequencing. Validation of disease-linked changes using GAPDH for normalization was observed for 8 out of the 12 circRNA studied in the cortex, and 5 out of 12 samples studied in the hippocampus (for the hippocampus a “trend level of significance” was used for validation because of the smaller sample size) (Figure 6A-B). circKATNAL2(hsa_circ_0108513) was significantly downregulated in both regions (Figure 6A-B). circKCNN2(hsa_circ_0127664), circPSMB1(hsa_circ_0078784), circMAPK9(hsa_circ_0001566) showed significant downregulation in the cortex, and a trend level decrease in the hippocampus samples. Multiple other circRNA showed differences in cortical samples that were significant and validated however did not show validation in the hippocampus (likely because of smaller sample sizes); these circRNA included: circRIMS1 (hsa_circ_0132250), circRAPGEF5(hsa_circ_0001681), circHOMER1(hsa_circ_0006916), circATRNL1(hsa_circ_0020093), circPDE4B(hsa_circ_0008433). qPCR plots normalized to circN4BP2L2 also gave similar results (Supplement Figure 5).

**Figure 6:**
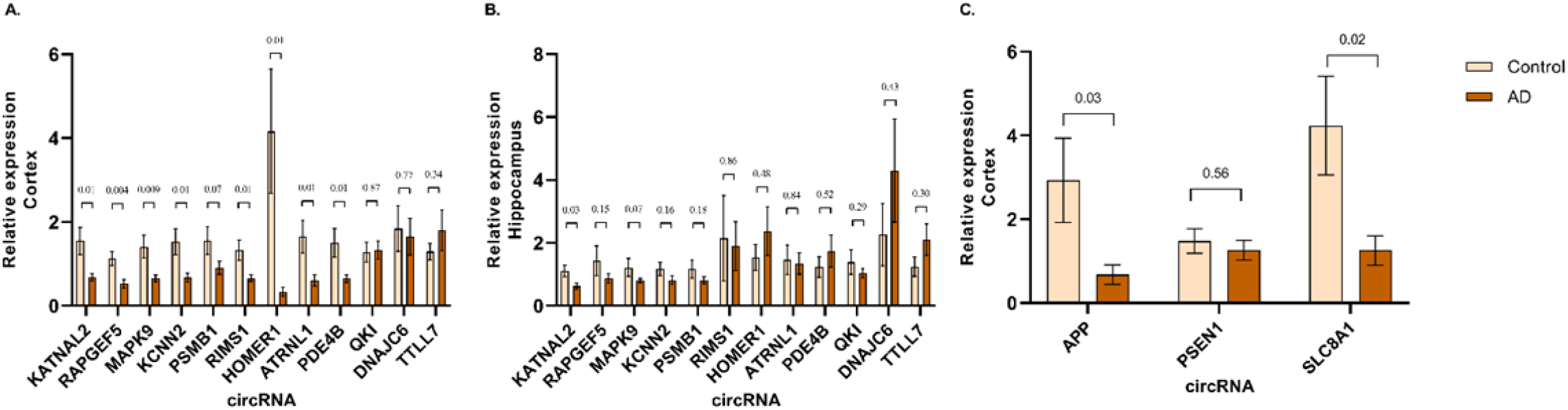
circRNA expression validation by RT-qPCR. (A and B) Relative expression of 12 circRNA checked in the cortex (N=20 AD, 20 Ctrl) and hippocampus (N=18 AD, 8 Ctrl), (C) Relative expression of circRNA associated with AD loci in the cortex. Data are given as mean ± SE.

We also used qPCR to validate circRNA produced from AD-associated genes, using cortical samples as a basis for the validation studies. The circAPP (hsa_circ_0003038) showed significant downregulation using either the GAPDH or circN4BP2L2 for normalization (Figure 6C, Supplement Figure 5D). The circPSEN1 (hsa_circ_0003848) also showed significant disease-linked increases expression using circN4BP2L2 but not GAPDH for normalization (Supplementary Figure 5D). Finally, we also examined the expression of circSLC8A1 which has been linked to Parkinson’s disorder ^19^. circSLC8A1 exhibited downregulation in the MSBB BM44 dataset (log FC=-0.27, p=0.004). qPCR of circSLC8A1 showed a strong and significant downregulation in the cortex of AD cases.

Of note, among the circRNA validated with qPCR, circATRNL1 is significantly associated with all three pathological traits. circHOMER1 and circPSMB1 are associated with CDR and Braak. circPSEN1 is significantly associated with plaque, however interestingly we find its expression is restricted to cortex. For Braak pathology, a correlation with circKCNN2 was observed.

### Neuronal exposure to oligomeric tau recapitulates disease-linked changed in circRNAs

Next, we investigated whether the changes in circRNA could be recapitulated in cell culture systems. We examined expression of the validated circRNA in neuronal precursors cells (NPCs) differentiated from human iPSCs. Next, we extracted oligomeric tau (oTau, S1p fraction) from the brain of PS19 P301S 9month aged mice, as described previously ^20-22^. The NPCs were treated with 80ng oligomeric tau (oTau) and harvested after 48 hrs, as described previously^21,22^, and qPCR was performed. As expected, oTau treatment of NPCs caused cytotoxicity (Supplementary figure 6A). The response of the NPCs to oTau was specific to some of the AD associated targets. oTau treatment decreased expression of circAPP, circPDE4B, circRAPGEF5, circMAPK9, circRIMS1 and circATRNL1 (Figure 7A). Interestingly, circPSEN1 also showed a decrease in expression upon oTau treatment, despite being upregulated in AD cases. circN4BP2L2 was tested as a negative control. To understand whether the circRNA differential regulation is specific to oTau treatment or induced by other stress responses, we subjected NPCs to oxidative stress using sodium arsenite. Interestingly, we observed no change in expression of AD associated circRNA upon arsenite treatment (Supplementary Figure 6B), which demonstrates that some circRNA are selectively reduced in response to oTau treatment. Finally, we performed qPCR analysis on mRNA to investigate whether changes in circRNA induced upon oTau treatment correlate with changes in their cognate linear transcripts. The response of most circRNAs examined (5/7) paralleled the response of the cognate mRNAs (Figure 7B). However, circMAPK9 exhibited downregulation independent of its linear transcript; oTau induced a decrease in circMAPK9 but not linear MAPK9 mRNA (Figure 7A, B). circAPP and circPSEN1 were more responsive to oTau than linear *APP* and *PSEN1* mRNA (Figure 7A, B). This highlights that circMAPK9, circAPP and circPSEN1 regulation in oTau treatment is specific.

**Figure 7:**
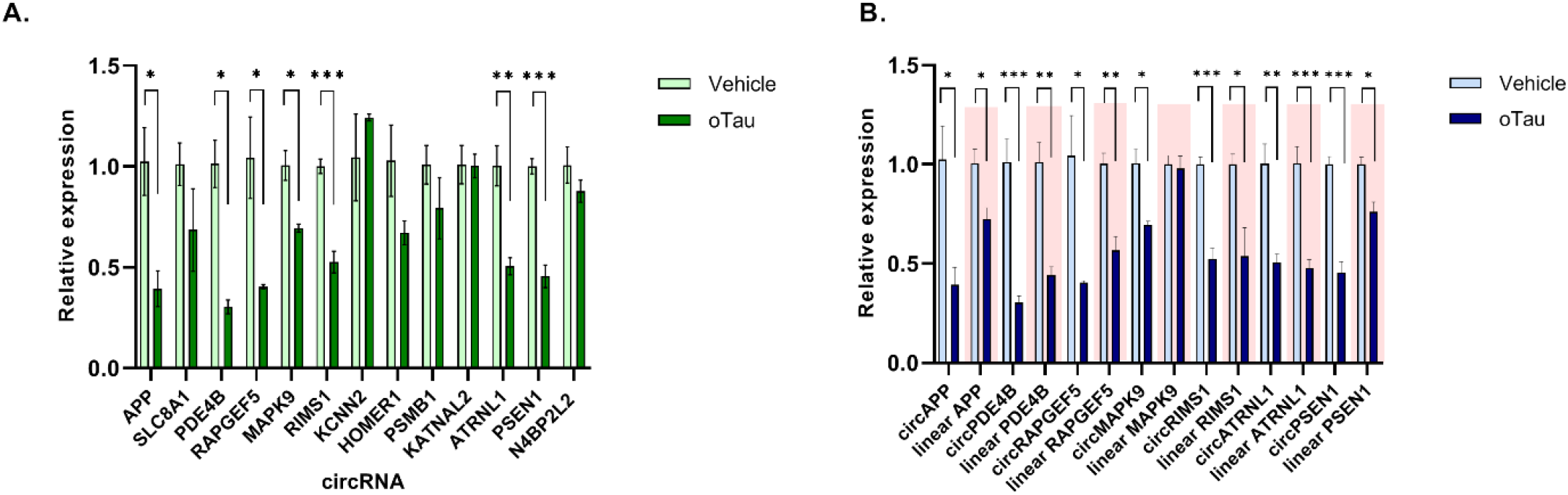
circRNA expression upon oTau treatment. (A) Relative expression of AD associated circRNA in NPCs with vehicle and oTau treatment. (B) Relative expression of cognate linear transcripts of circRNA modulated with tau toxicity in NPCs. Data are given as mean ± SE, n = 3 biological replicates. *p<0.05, **p<0.01, ***p<0.001

## Discussion

CircRNAs comprise an important segment of the transcriptome that is particularly abundant in brain, yet its contributions to neurodegenerative diseases are poorly understood. The pipeline that we established enables exploration of circRNA expression to identify those circRNA exhibiting the strongest changes with disease.

CircRNAs are particularly interesting because they localize to 2 organelles critical to the disease process, synapses and stress granules (the latter being a membraneless organelle) ^23,24^. Dysfunction of circRNA biology in either of these organelles could directly impact on the disease process. Indeed, a recent study of circRNAs identified in 5X-FAD mice identified a circRNA, Cwc27, that is increased with the disease. Conversely, knockdown of Cwc27 reduced Aβ accumulation by sequestering the RNA binding protein Pur-α in the cytoplasm, and (importantly) increasing the expression of neprilysin ^25^. The circRNAs that we observed are notable for the disease linkage and brain distribution. Our study employs the newest pipelines for circRNA identification, and is also the first to examine the hippocampus, which is the region most affected in AD. The circRNAs that we identified showing the strongest DE linked to AD exhibited patterns of DE expression that correlated with indices of tauopathy (Braak staging) and cognitive loss (CDR) suggesting that these circRNAs reflect neuronal biology, which is consistent with the current understanding of circRNA functions. Interestingly, the DE circRNAs that we observed did not show DE for the corresponding mRNA, which raises the possibility that the biology of circRNA either more directly contributes to or is more sensitive to the pathophysiology of AD.

In order to understand the regulation of disease linked circRNAs, we examined circRNA expression in NPCs. The first observation of note was the higher level of expression of the circRNAs in human NPCs versus HEK 293 cells, which we speculate reflects the neurobiology of neurons. The increase in circRNA expression occurring with differentiation of NPCs supports a hypothesis that neuronal phenotypes lead to higher circRNA expression. The downregulation of the validated circRNAs following treatment with oTau highlights the responsiveness of circRNAs in NPC to regulation, and suggests that oTau elicits regulation of circRNAs that is similar between NPCs and the brain. These observations also raise the possibility that some of the changes in circRNA expression in the AD brain might reflect the response to oTau toxicity. This observation would be consistent with Cervera-Carles et al who noted similar responses of circRNAs between FTD-tau (but not FTD-TDP-43) and AD^15^. Future studies will now be able to explore the mechanisms through which oTau regulates circRNA production.

Strengths and Limitations: The strengths of this study derive from the application of the latest algorithms to the study of human brain, the examination of multiple brain regions and the extension of this work to NPCs, which provides a potential model for studying the regulation of circRNAs. However, the study also has important limitations. The small number of hippocampal samples limited the power of these investigations, with a resulting reduced ability to detect significant disease-linked changes using hippocampal samples. Another weakness is the correlational nature of the studies. Future studies will need to focus on modulating levels of circRNA to determine whether circRNA species can impact on the pathophysiology of AD.

## Conclusion

CircRNAs are a novel class of regulatory RNAs but their role in AD pathogenesis is just beginning to be explored. Our study identified circRNAs whose expression is linked to AD and are correlated with indices of tauopathy and cognitive loss in the cortex and hippocampus. Furthermore, particular circRNAs were differentially expressed among patients with AD or other dementias, suggesting that circRNA might be useful disease biomarkers or targets for disease intervention.

## Acknowledgements

Brain tissues for this study were provided by the NIH NeuroBioBank, the Boston University Alzheimer’s disease center (NIH grants AG50204517 and P30-AG13846) and the Goizueta Alzheimer’s Disease Research Center (ADRC) of Emory University (P30 AG066511). Funding for this grant was provided by NIH: AG0772577 BW/XZ, AG056318 (BW), AG074591 (BW), AG061706 (BW), AG064932 (BW) and the BrightFocus Foundation (BW).

## Methods

### Generating and Quality Control of the BU-ADRC Hippocampus RNA-seq data

Frozen human brain hippocampal tissues were collected at the BU-ADRC. After RNA extraction, RNA was purified using RNeasy Mini Kits (Catalog No. 74134, Qiagen, Germany). The RNA integrity number was calculated using an RNA 6000 Pico assay on a Bioanalyzer 2100 (Agilent Technologies, USA). The extracted RNA was also quantified using the Quant-iT RNA assay (Catalogue no. Q33140, Invitrogen, USA) on a Qubit Fluorometer (Thermo Fisher Scientific). Selected high-quality RNA were shipped to the Genome Center at Yale University in New Haven. Total RNA quality was determined by estimating the A260/A280 and A260/A230 ratios on nanodrop. RNA integrity was determined by Agilent Bioanalyzer. rRNA was depleted using the Kapa RNA HyperPrep Kit with RiboErase (Catalog No. KR1351, Kapa Biosystems, USA). Following sample fragmentation, adapter ligation, and library amplification, 100bp paired-end strand-specific sequencing was performed on an Illumina NovaSeq according to Illumina protocols. All samples were randomly assigned to a sequencing pool before sequencing, and RNA extraction and sequencing library preparation were performed blind to neuropathological case–control status. The average number of raw sequencing reads per individual was 40M (details of RNA quality control, RNA-seq library preparation, and sequencing are in Supplementary Methods).

We received 222 paired FASTQ files for 211 subjects of our BU-ADRC hippocampus samples. All datasets were subjected to three levels of quality check: 1) FastQC was used to inspect the initial fastq files that visually reflect sequencing quality and mismatching of sex (http://www.bioinformatics.babraham.ac.uk/projects/fastqc/). Low-quality reads (5% of the total) and poor-quality base were eliminated and adaptor sequences were trimmed using Trimmomatic (version 3.9)^26^. which is a fast, multithreaded command-line tool for trimming, cropping, and removing adapters from Illumina fastq data. 2)The percentage of mapped and unmapped reads was evaluated. 3) Qualification analysis of the aligned results was performed by a) checking the library size and gene expression level distribution for sample quality check and duplicate samples detection; b) checking sex marker gene expression level across all samples; c) PCA analysis for identifying hidden batch effects. One sample was identified as a duplicate and was excluded.

### Preprocessing two public AD brain RNA-seq datasets

The Mount Sinai Brain Bank (MSBB) cortex RNA-seq dataset was downloaded from the Synapse portal (syn315774). In short, four different cortical regions were generated, which are the frontal pole (Brodman area (BM10), superior temporal gyrus (BM22), parahippocampal gyrus (BM36), and inferior frontal gyrus tissue (BM44)) from 301 individuals. The Ribo-Zero rRNA Removal Kit (Illumina human/mouse/rat) was applied to remove the rRNAs. The sequencing libraries preparation used TruSeq RNA Sample Preparation kit v.2. Using the Illumina HiSeq 2500, rRNA-depleted, 101-nt, single-end, and non-stranded RNA-seq data were obtained from these libraries. In this study, only MSBB BM44 cortex data were analyzed.

The Adult Changes in Thought (ACT) hippocampus RNA-seq dataset was downloaded from NIAGADS (NG00059, https://www.niagads.org/datasets/ng00059), which contains 25 AD cases and 50 normal controls. After rRNA-depletion, 51bp paired-end and stranded RNA-seq data were generated using the Illumina HiSeq 2500. After downloading the raw fastq files, the same preprocessing and quality check pipeline used above for our BU-ADRC Hippocampus RNA-seq data was applied to these two public AD brain RNA-seq datasets.

### CircRNA detection and quantification using CIRCexplorer3-CLEAR

To detect circRNA from rRNA-depleted RNA-seq data, we implemented two different pipelines as shown in **Figure 1**. CIRCexplorer3-CLEAR is a pipeline designed for comparing the expression of circular and linear RNAs^27^, which includes two main steps: alignment and quantification. The trimmed RNA-seq reads were firstly aligned to the human reference genome (GRCh38/hg38) using HISAT2 (version 2.0.5)^28^ with known gene annotations for subsequent linear RNA quantification. HISAT2-mapped fragments were used to quantify and select the maximally-expressed transcript of a given gene using StringTie (version 1.3.3). FPB (fragments per billion mapped base) value was applied to quantitate/normalize linear RNA expression by HISAT2-mapped fragments to splicing-junction (SJ) sites of the maximally-expressed transcript annotation.

Separately, HISAT2-unmapped fragments were mapped to the same hg38 reference genome using TopHat-Fusion (version 2.0.12) ^29^ for subsequent circRNA quantification. Then, fragments mapped to back-splicing-junction (BSJ) sites were retrieved from TopHat-Fusion (version 2.3.6; *CIRCexplorer2 parse -f -t TopHat-Fusion*) ^30^ and normalized by totally mapped bases to obtain FPB values for circRNA quantification. circRNA were called for each sample separately and were manually combined as a data matrix for further analysis.

*clear_quant -g hg38*.*fa \ -i hisat_index \ -j bowtie1_index \ -G hg38_ref_all2*.*gtf*

Finally, the number of detected circRNAs were summarized by applying different cutoff, for example, FPB> 0, 0.1, 0.2, 0.5, or 1.0 in over 25%, 50%, 70%, or 100% of total samples. circRNAs with FPB> 0 in over 50% of samples were selected for downstream differential expression analysis. For these selected circRNAs, the ratio of corresponding linear RNAs and circRNAs is great than 0.1 in at least 3 samples. Just like CIRCexplorer3, DCC has two main steps (details were presented in Supplementary Methods).

### Identifying circRNAs differentially expressed between AD and control in the hippocampus and cortex

CircRNAs that were not expressed in more than 50% of the samples were excluded from the analysis. Differential expression analyses for CIRCexplorer3 calling circRNA between AD cases and control were performed using Limma^31^. The FPB circ matrix with log2(x + 0.5) transformation was compared between AD cases and controls using linear regression models adjusting for sex, age, RQN (QNA quality number), and library batch.

Before running limma, CIRCexplorer3 circRNA counts in each sample were normalized by fragments per billion mapped bases (FPB). The normalized FPB values were taken log2(x + 0.5) transformation. Subjects with missing covariate data and disease subgroup information were excluded. Only circRNAs expressed in more than 50% of the samples were included in the DE analysis. In the model, technical differences (RQN/PMI, batch), age/ Age of Death (AOD), and sex were included as covariates.

Meta-analysis of hippocampus and BM44 differential results of AD vs. control was performed using METAL^32^ under a fixed effects model. A statistical significance threshold was defined as FDR<0.05/0.1. Significant circRNA-related cognate genes were used for pathway enrichment analysis using EnrichR ^33^.

### circRNA association analysis with AD neuropathology and related endophenotypes in hippocampus and cortex

In addition to traditional case-control analysis, we investigated the circRNA association with AD neuropathological stage including amyloid CERAD score, tau Braak score, and clinical dementia rate with each study. In the BU-ADRC hippocampus data, a low CERAD score means high amyloid burden and vice versa. However, in the MSBB BM44 cortex data, plaque mean (PM) value was provided with a high PM value indicates high amyloid burden. Therefore, in the meta-analysis of amyloid plaque burden, the direction of the coefficient estimated from BU-ADRC hippocampus data was first reversed, then input to METAL.

### circRNA differential expression analysis in AD and mixed AD in the hippocampus

For BU-ADRC hippocampus RNA-seq data, similar to circRNAs differential expression analysis between AD and normal controls, mixed AD with vascular dementia (VaD) or AD with Lewy Boby (LBD) were compared to control to discover unique and common circRNAs differential expression patterns with the same covariates included in the analytic model using limma.

### Quantitative real-time PCR (qRT-PCR)

30mg of human brain tissue was homogenized in 1ml of TRIzol (Catalog No. 15596026, Thermofisher Scientific, USA). 200µl of chloroform was added to each sample and mixed well. The samples were centrifuged at 13,000rpm for 15 minutes at 4°C. The RNA in the upper aqueous layer was subjected to column-based purification using the Qiagen RNeasy Plus mini kit (Catalog No. 74134, Qiagen, Germany). RNA was reversed transcribed using random hexamer primers with High-Capacity cDNA Reverse Transcription Kit (Catalog No. 4368814, Thermofisher Scientific, USA). For qPCR, a reaction mixture of 1µl cDNA, 0.2µM primers, and 2.5µl Ssoadvanced™ Universal SYBR (Catalog No. 1725720, Bio-Rad, USA) upto 5µl was prepared in triplicates for each sample. qPCR was performed in Quantstudio 12K Flex qPCR System (Thermofisher Scientific, USA). Divergent primers were used to amplify circRNA. However, in some instances the primers amplified more than one isoform for the circRNA and hence specific isoforms were amplified using primers at back-splice junction. The sequence for primers used for amplification are provided in Supplementary Table. The relative expression levels were quantified with 2^−ΔΔCt^ method and either GAPDH or circN4BP2L2 was used for normalization. For calculating statistical significance unpaired t-test was used. All statistical analysis was done using GraphPad Prism 9.

### Cell culture treatments

Neuronal precursor cells (NPCs) were obtained from ATCC (Catalog no. ACS 5004, ATCC, USA). The cells were maintained in ATCC-NPC medium (Catalog no. ACS 3003, ATCC, USA) in a 37°C incubator containing 5% CO2. oTau oligomers were generated from the brain of 9-month-old PS19 P301S mice by the fractionation method described previously ^20-22^. 60,000 cells were plated in each well of 24 well plate in triplicates for each condition. NPCs were treated with 80ng of oTau oligomers for 48hrs. Post-treatment, RNA was isolated from cells and the spent media was analyzed for cytotoxicity using CyQUANT LDH Cytotoxicity Assay Kit(Catalog no. C20300, Invitrogen, USA). For oxidative stress, NPCs were treated with 500µM of sodium arsenite for 1 hour.

