## Supplementary Information for "Identification of circRNAs linked to Alzheimer’s disease and related dementias"

**Supplemental Materials/Online Materials**

**Supplemental Methods**

**RNA quality control, RNA-seq library preparation, and sequencing of BU-ADRC hippocampus samples at the Genome Center at Yale University**

RNA Seq Quality Control: Total RNA quality is determined by estimating the A260/A280 and A260/A230 ratios by nanodrop. RNA integrity is determined by running an Agilent Bioanalyzer gel, which measures the ratio of the ribosomal peaks. Samples with RNA integrity number (RIN) values of 5 or greater are recommended for library preparation.

RNA Seq Library Preparation: Using the Kapa RNA HyperPrep Kit with RiboErase (Catalog No. KR1351, Kapa Biosystems, USA), rRNA is depleted starting from 25-1000ng of total RNA by hybridization of rRNA to complementary DNA oligonucleotides, followed by treatment with RNase H and DNase to remove rRNA duplexed to DNA. Samples are then fragmented using heat and magnesium. 1^st^ strand synthesis is performed using random priming. 2^nd^ strand synthesis incorporates dUTPs into the 2^nd^ strand cDNA. Adapters are then ligated and the library is amplified. Strands marked with dUTPs are not amplified allowing for strand-specific sequencing. Indexed libraries that meet appropriate cut-offs for both quantity and quality are quantified by qRT-PCR using a commercially available kit (KAPA Biosystems, USA) and insert size distribution determined with the LabChip GX or Agilent Bioanalyzer. Samples with a yield of ≥0.5 ng/ul are used for sequencing.

Flow Cell Preparation and Sequencing: Sample concentrations are normalized to 1.2 nM and loaded onto an Illumina NovaSeq flow cell at a concentration that yields 25 million passing filter clusters per sample. Samples are sequenced using 100bp paired-end sequencing on an Illumina NovaSeq according to Illumina protocols. The 10bp unique dual index is read during additional sequencing reads that automatically follow the completion of read 1. Data generated during sequencing runs are simultaneously transferred to the YCGA high-performance computing cluster. A positive control (prepared bacteriophage Phi X library) provided by Illumina is spiked into every lane at a concentration of 0.3% to monitor sequencing quality in real-time.

Data Analysis and Storage: Signal intensities are converted to individual base calls during a run using the system's Real-Time Analysis (RTA) software. Base calls are transferred from the machine's dedicated personal computer to the Yale High-Performance Computing cluster via a 1 Gigabit network mount for downstream analysis. Primary analysis - sample de-multiplexing and alignment to the human genome - is performed using Illumina's CASAVA 1.8.2 software suite. The data are returned to the user if the sample error rate is less than 2% and the distribution of reads per sample in a lane is within a reasonable tolerance. Data is retained on the cluster for at least 6 months, after which it is transferred to a tape backup system.

**CircRNA alignment and quantification using DCC pipeline**

For the DCC pipeline, the chimeric read was generated using chimeric read-detection mode in STAR, which aligned the reads from both MSBB RNA-seq datasets to the GENCODE28-annotated human reference genome (GRCh38). Then chimeric reads were further processed and filtered using DCC software to identify backsplice junctions. Finally, the backsplice junction numbers were compressed on to their linear gene of origin to obtain a collection of high-confidence circRNA counts for downstream studies.

To detect circRNA from rRNA-depleted RNA-seq data circRNA, we implemented two different pipelines as shown in **Supplementary** **Figure 1**.

A. RNA-seq data preprocessing and alignment

We used STAR (version 2.6.1c) ^[1]^ with chimeric read-detection mode and suggested parameters in circRNA calling software DCC to align the reads from RNA-seq datasets. In the beginning, we created genomics alignment index files with a splice junction database with a read length of 100 using GRCh38.95.gtf as reference annotation. For each paired FASTQ file, we did individually and together alignment with the following parameters:--readFilesCommand zcat\ --outSAMtype BAM SortedByCoordinate \ --outFileNamePrefix ${star_out_dir}/mate2/${read_2}_ \ --outSJfilterOverhangMin 15 15 15 15 \ --alignSJoverhangMin 15 \ --alignSJDBoverhangMin 15 --seedSearchStartLmax 30 --outFilterMultimapNmax 20 --outFilterScoreMin 1 \ --outFilterMatchNmin 1 --outFilterMismatchNmax 2 --chimSegmentMin 15 --chimScoreMin 15 \ --chimScoreSeparation 10 --chimJunctionOverhangMin 15 \ --genomeLoad LoadAndKeep --limitBAMsortRAM 30000000000 \

B. CircRNA detection from back-splicing region

DCC is a software tandem that systematically identifies, quantifies, and filters circRNA from chimeric junctions generated during STAR alignment ^[2]^. We followed the recommended parameters for filtering from DCC: 1) All backsplice junctions found in repetitive regions of the genome were excluded. 2) all reads from the mitochondrial chromosome and spanned multiple gene annotations were skipped. For better accuracy, we put both mates individually and together when we performed DCC software.

DCC -D -R \hg_simple_repeat.gtf -an gencode.v26.primary_assembly.annotation.gtf -Pi -F -M -Nr 5 10 \-fg -G -A GRCh38.primary_assembly.genome.fa

**Differential expression analysis for circRNA detected from DCC**

Besides using limma for circRNA normalized FPB matrix by Circexplore3, we also performed differential expression analysis for circRNA raw count matrix built from DCC using DEseq2. The raw count from DCC was normalized on the basis of a sequencing depth/library-size-derived size factor in DEseq2. The same covariates technical differences (RQN/PMI, batch), age/ Age of Death (AOD), and sex were included in the analytic model.

**RNA-seq alignment and quantification of linear mRNA:** The hippocampus RNA-seq data was aligned to the human reference genome (GRCh38.95) using STAR (version 2.6.1c) as read aligners which use 2-pass mapping to improve the mapping chances of splice reads from novel junctions^[1,3]^. For reading of trimmed FASTQ files, the *readFilesCommand* option for reading mapped reads has been used, and *GeneCounts* for counting mapped reads per gene under the *quantMode* mapping mode and then *twoPassMode* options. *TranscriptomeSAM* mode was applied to translate transcript coordinates option for mapped reads. For each sample, a BAM file with a corresponding alignment report file was generated.

The RSEM (version 1.3.1) ^[4]^ and Bowtie2 (version 2.3.4.1) ^[5]^ were used for gene and isoform levels quantification and Homo sapiens.GRCh38.95.gtf was implemented as annotation files. For each sample, this process generated gene and isoform expression data that included the gene id, gene length, effective gene length, expected count, counts per million (CPM), and FPKM reads.

**Supplementary Table 1.** Mount Sinai Brain Bank (MSBB) RNAseq data downloaded from Accelerating Medicines Partnership–Alzheimer’s Disease (AMP-AD) knowledge portal.

| **Distribution of subjects in MSBB cortex BM44 data (N=308)** | | | | | |
| --- | --- | --- | --- | --- | --- |
| **Variable** | **AD** | **Control** | **Variable** | **AD** | **Control** |
| **N** | 216 | 92 | **Braak** |  |  |
| **Age(yr)**  **(SD)** | 84.2  (6.85) | 79.9  (8.76) | **0** | 1 | 10 |
| **Female N**  **(%)** | 148  (68.52) | 45  (48.91) | **1** | 8 | 18 |
| **PMI** |  |  | **2** | 12 | 30 |
| **< 10** | 182 | 55 | **3** | 35 | 28 |
| **10-19.9** | 31 | 26 | **4** | 30 | 2 |
| **20-29.9** | 3 | 10 | **5** | 29 | 1 |
| **>= 30** | 0 | 1 | **6** | 95 | 0 |
| **CDR** | | | **Plaque Mean** | | |
| **0** | 14 | 30 | **< 2** | 13 | 80 |
| **0.5** | 16 | 32 | **2-6.9** | 66 | 11 |
| **1** | 20 | 10 | **7-9.9** | 51 | 1 |
| **2** | 37 | 7 | **>= 10** | 86 | 0 |
| **3** | 66 | 6 |  |  |  |
| **4** | 30 | 6 |  |  |  |
| **5** | 33 | 1 |  |  |  |

**Supplementary Table 2.** the Adult Changes in Thought (ACT) RNAseq data were downloaded from the NIA Genetics of Alzheimer's Disease Data Storage Site (NIAGADS).

| **Distribution of subjects in ACT Hippocampus data (N=75**) | | |
| --- | --- | --- |
| **Variable** | **AD** | **Control** |
| **N** | 25 | 50 |
| **Age(yr) (SD)** | 91.16 (7.05) | 89.06 (6.89) |
| **Female** | 52% | 40% |
| **Race** |  |  |
| **1** | 0 | 2 |
| **2** | 25 | 48 |
| **Braak** | | |
| **0** | 1 | 2 |
| **1** | 1 | 10 |
| **2** | 2 | 9 |
| **3** | 2 | 15 |
| **4** | 4 | 7 |
| **5** | 7 | 5 |
| **6** | 8 | 2 |
| **CERAD** | | |
| **0** | 5 | 12 |
| **1** | 2 | 21 |
| **2** | 7 | 12 |
| **3** | 11 | 5 |

**Supplementary Table 3**. CircRNA detected in hippocampus and cortex regions

|  | **Cortex (MSBB BM44)** | | | **Hippocampus (BU-ADRC)** | | |
| --- | --- | --- | --- | --- | --- | --- |
| **Number of samples expressed** | **#circRNAs** | **#unique gene** | **#circRNA**  **per gene** | **#circRNAs** | **#unique gene** | **#circRNA**  **per gene** |
| FPB > 0 and over 25% of sample expressed | 10,988 | 3,912 | 2.81 | 9,604 | 3,523 | 2.73 |
| **FPB > 0 and over 50% of sample expressed** | **4,912** | **2,414** | **2.03** | **4,092** | **2,056** | **1.99** |
| FPB > 0 and over 75% of sample expressed | 2,212 | 1,359 | 1.63 | 1,762 | 1,125 | 1.57 |
| FPB > 0 and over 100% of sample expressed | 84 | 75 | 1.12 | 0 | 0 | NA |
| FPB > 0.1 and over 25% of sample expressed | 10,988 | 3,912 | 2.81 | 7,945 | 3,176 | 2.50 |
| FPB > 0.1 and over 50% of sample expressed | 4,912 | **2,414** | 2.03 | 3,570 | 1,865 | 1.91 |
| FPB > 0.1 and over 75% of sample expressed | 2,212 | 1,359 | 1.63 | 1,597 | 1,038 | 1.54 |
| FPB > 0.1 and over 100% of sample expressed | 84 | 75 | 1.12 | 0 | 0 | NA |
| FPB > 0.5 and over 25% of sample expressed | 5,741 | 2,660 | 2.16 | 2,095 | 1,268 | 1.65 |
| FPB > 0.5 and over 50% of sample expressed | 3,018 | 1,694 | 1.78 | 1,173 | 799 | 1.47 |
| FPB > 0.5 and over 75% of sample expressed | 1,583 | 1,026 | 1.54 | 620 | 456 | 1.36 |
| FPB > 0.5 and over 100% of sample expressed | 48 | 47 | 1.02 | 0 | 0 | NA |
| FPB > 0.8 and over 25% of sample expressed | 3,765 | 1,989 | 1.89 | 1,242 | 831 | 1.49 |
| FPB > 0.8 and over 50% of sample expressed | 2,111 | 1,310 | 1.61 | 702 | 507 | 1.38 |
| FPB > 0.8 and over 75% of sample expressed | 1,203 | 803 | 1.50 | 377 | 299 | 1.26 |
| FPB > 0.8 and over 100% of sample expressed | 27 | 26 | 1.04 | 0 | 0 | NA |
| FPB > 1 and over 25% of sample expressed | 2,907 | 1,659 | 1.75 | 942 | 667 | 1.41 |
| FPB > 1 and over 50% of sample expressed | 1,705 | 1,095 | 1.56 | 543 | 407 | 1.33 |
| **FPB > 1 and over 75% of sample expressed** | **980** | **677** | **1.45** | **285** | **232** | **1.23** |
| FPB > 1 and over 100% of sample expressed | 24 | 23 | 1.04 | 0 | 0 |  |
| FPB > 1.5 and over 25% of sample expressed | 1,879 | 1,195 | 1.57 | 555 | 419 | 1.32 |
| FPB > 1.5 and over 50% of sample expressed | 1,164 | 780 | 1.49 | 323 | 257 | 1.26 |
| FPB > 1.5 and over 75% of sample expressed | 677 | 510 | 1.33 | 168 | 141 | 1.19 |
| FPB > 1.5 and over 100% of sample expressed | 6 | 6 | 1.00 | 0 | 0 | NA |
| FPB > 2 and over 25% of sample expressed | 1,368 | 902 | 1.52 | 369 | 292 | 1.26 |
| FPB > 2 and over 50% of sample expressed | 838 | 602 | 1.39 | 205 | 169 | 1.21 |
| **FPB > 2 and over 75% of sample expressed** | **512** | **400** | **1.28** | **114** | **103** | **1.11** |
| FPB > 2 and over 100% of sample expressed | 3 | 3 | 1.00 | 0 | 0 | NA |

**Supplemental Tables 4.** circRNAs expressed at >= 1 FPB in over 75% samples of hippocampus (**see Excel**)

**Supplemental Tables 5.** circRNAs expressed at >= 1 FPB in over 75% samples of cortex (**see Excel**)

**Supplemental Tables 6.** DE circRNAs (n=218 at FDR<0.1) in an independent ACT hippocampus dataset (**see Excel**)

**Supplemental Tables 7.** Gene ontology analysis of gene loci with 218 circRNAs differentially expressed between AD and controls (**see Excel**)

**Supplementary Table 8.** meta-analysis of 218 circRNA associated with AD pathology and CDR in hippocampus and cortex (**see Excel**)

**Supplemental Tables 9.** Details for samples used for qPCR validation: cortex (**see Excel**)

**Supplemental Tables 10.** Details for samples used for qPCR validation: hippocampus (**see Excel**)

**Supplemental Tables 11.** Primers used for qPCR validation (**see Excel**)

**Supplementary Figure 1**. **Analysis pipeline of circRNA alignment and quantification in AD brains.** (A) Flowchart for the pipeline for circRNA calling. The hippocampus data and MSBB cortex region were used for analysis. Both DCC and circexplore3 were used for calling but we show only Circexplore3 results in the main figure and tables. (B) Pipeline for circRNA calling using DCC. (C) Pipeline for circRNA calling using Circexplore3.

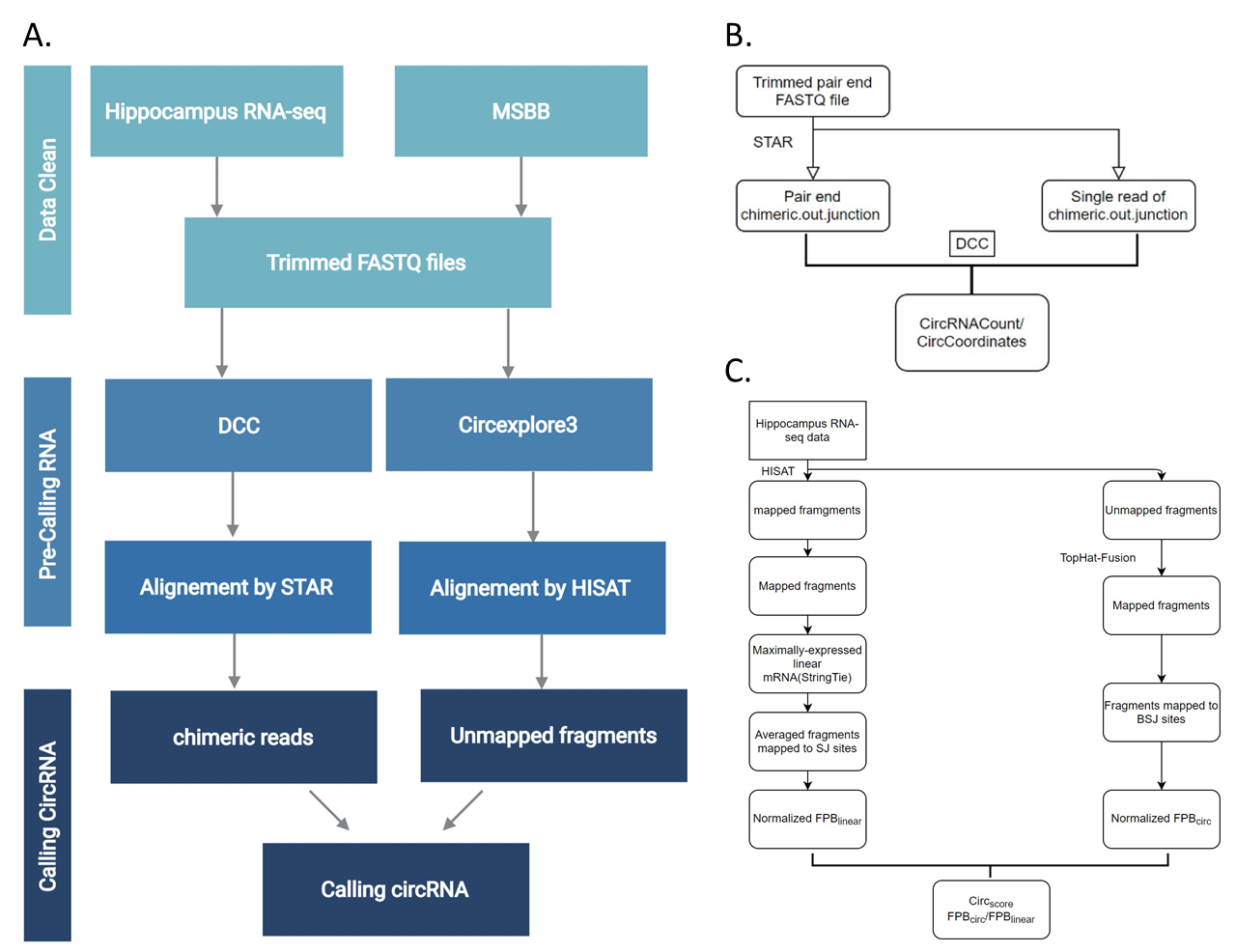

**Supplementary Figure 2**. Gene ontology analysis of gene loci with circRNAs expressed at FPB>1 (FPB>2) in over 75% samples in hippocampus and cortex.

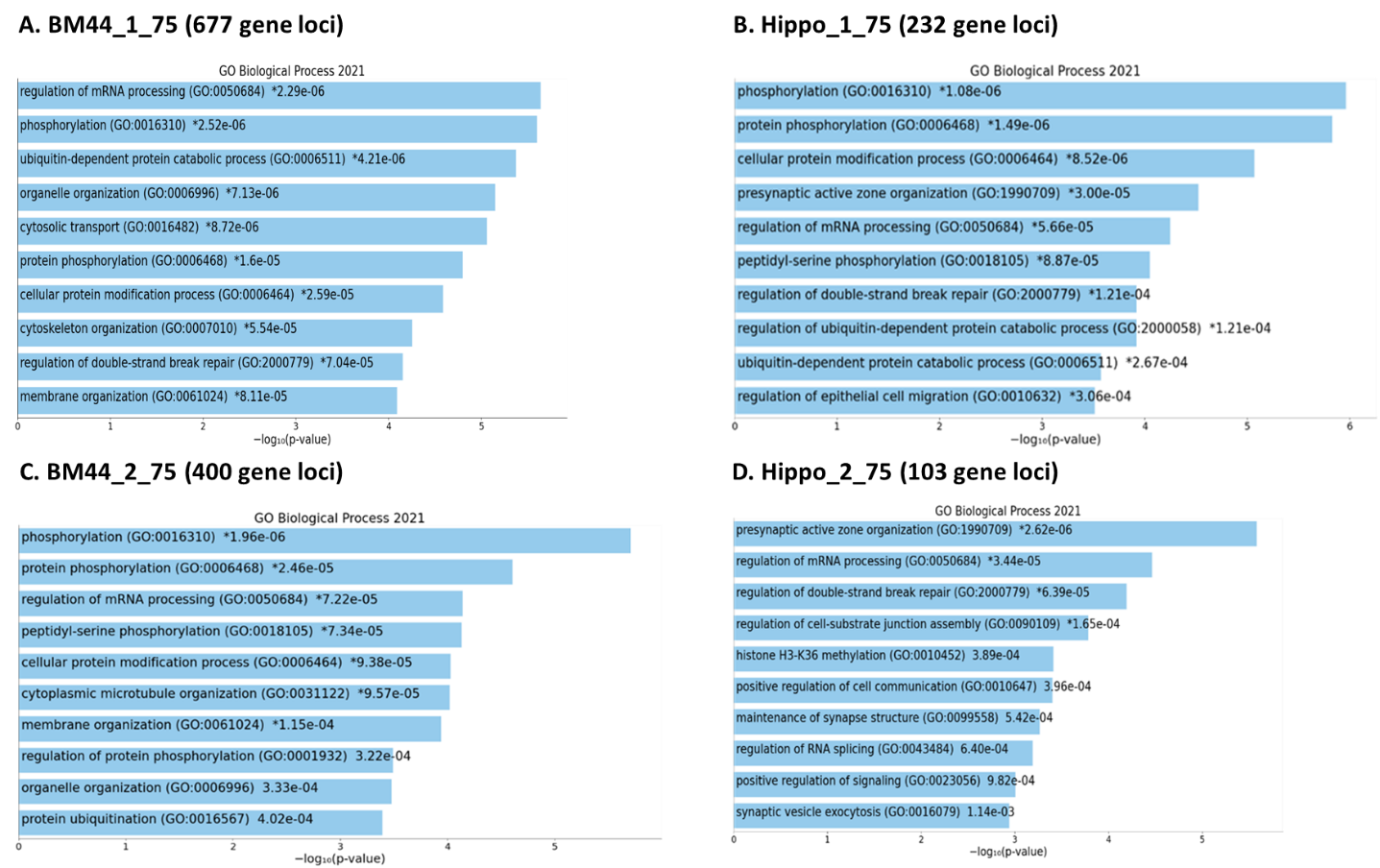

**Supplementary Figure 3**. Heatmap of 48/218 circRNAs in the hippocampus of control, pure AD, AD+LBD and AD+VaD.

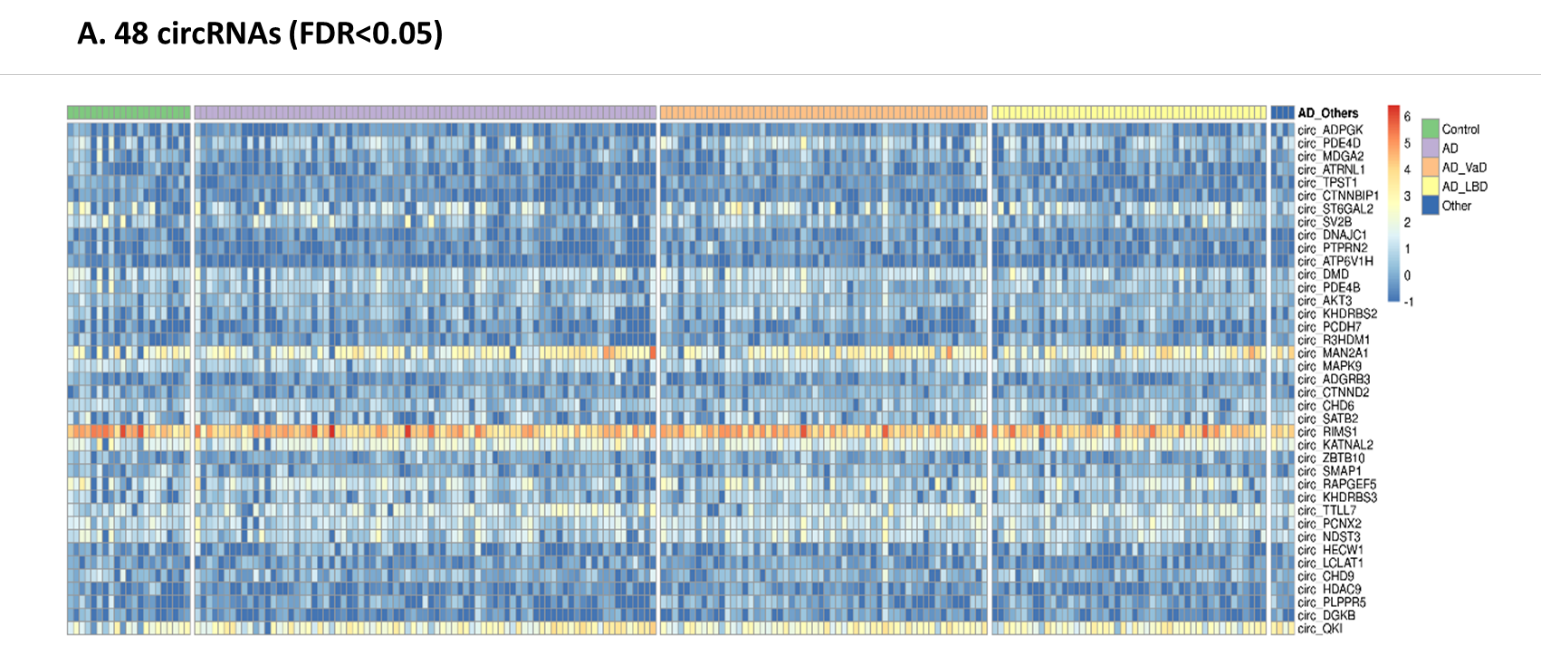

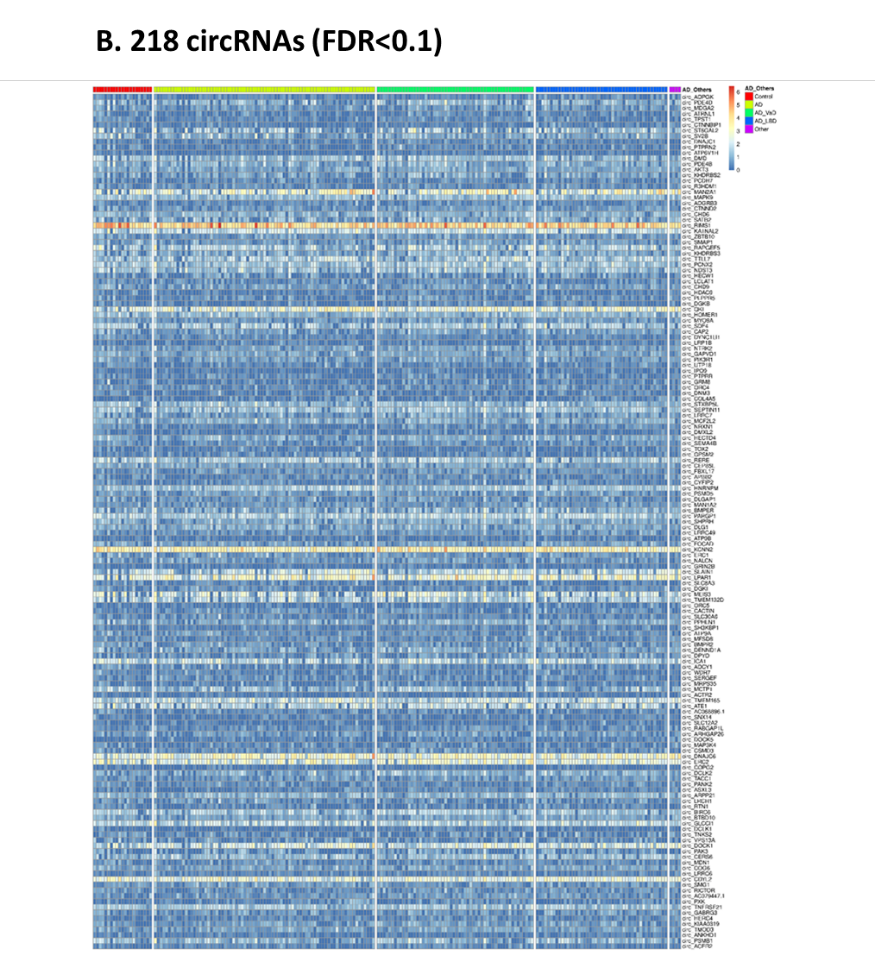

**Supplementary Figure 4.** Venn Diagram of 218 DE circRNAs associated with AD pathology and CDR in hippocampus and cortex.

| **218 circRNAs (AD vs. Control at FDR<0.1)** | |
| --- | --- |
| 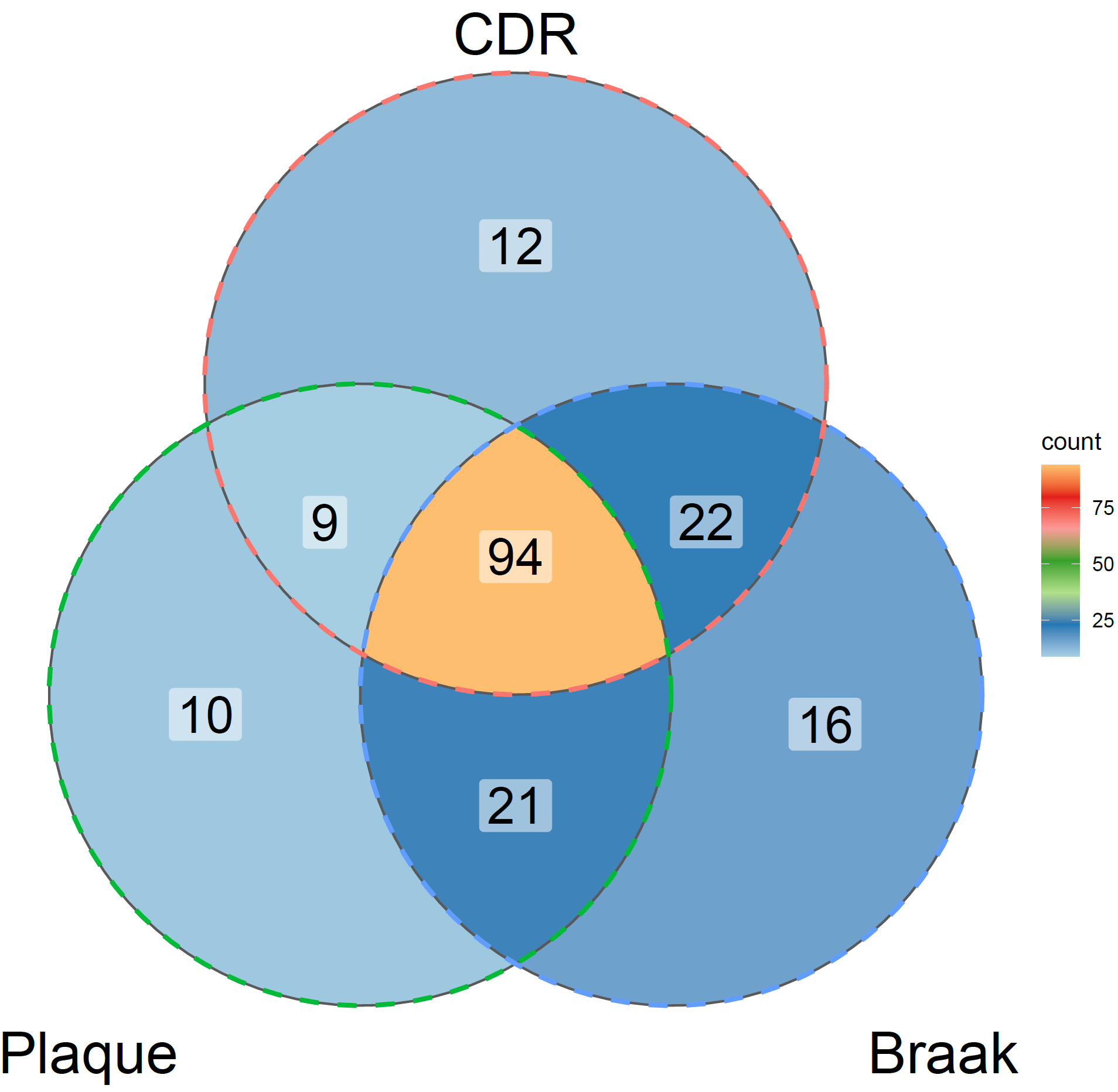 | 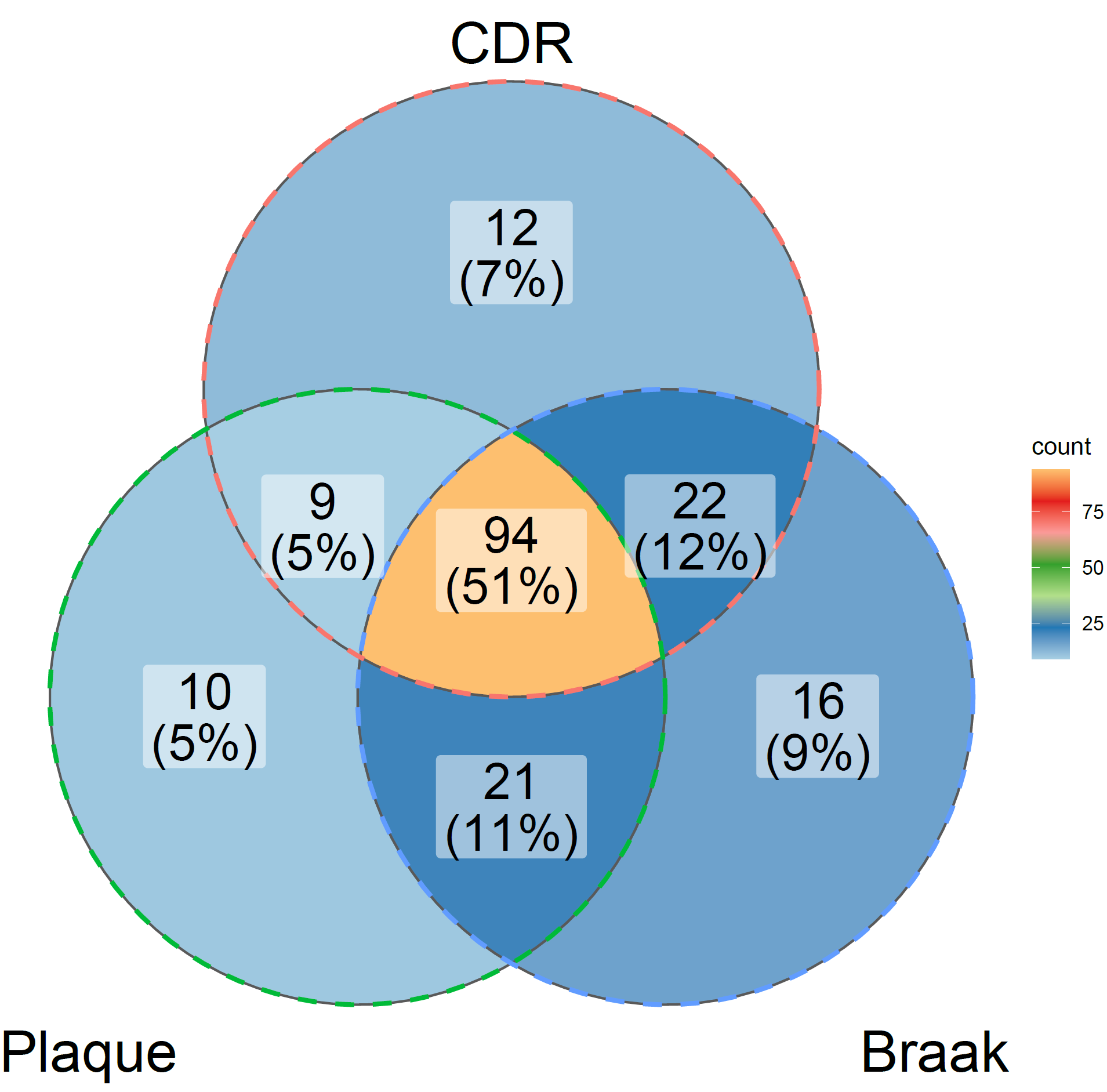 |

**Supplementary Figure 5**:  **circRNA expression validation by RT-qPCR using circN4BP2L2 as the internal control.** (A) No change in the relative expression of circN4BP2L2 in AD compared to controls, normalized to GAPDH. (B and C) Relative expression of 12 circRNA checked in the cortex (N=20 AD, 20 Ctrl) and hippocampus (N=18 AD, 8 Ctrl), normalized to circN4BP2L2. (D) Relative expression of circRNA associated with AD loci in the cortex, normalized to circN4BP2L2. Data are given as mean ± SE.

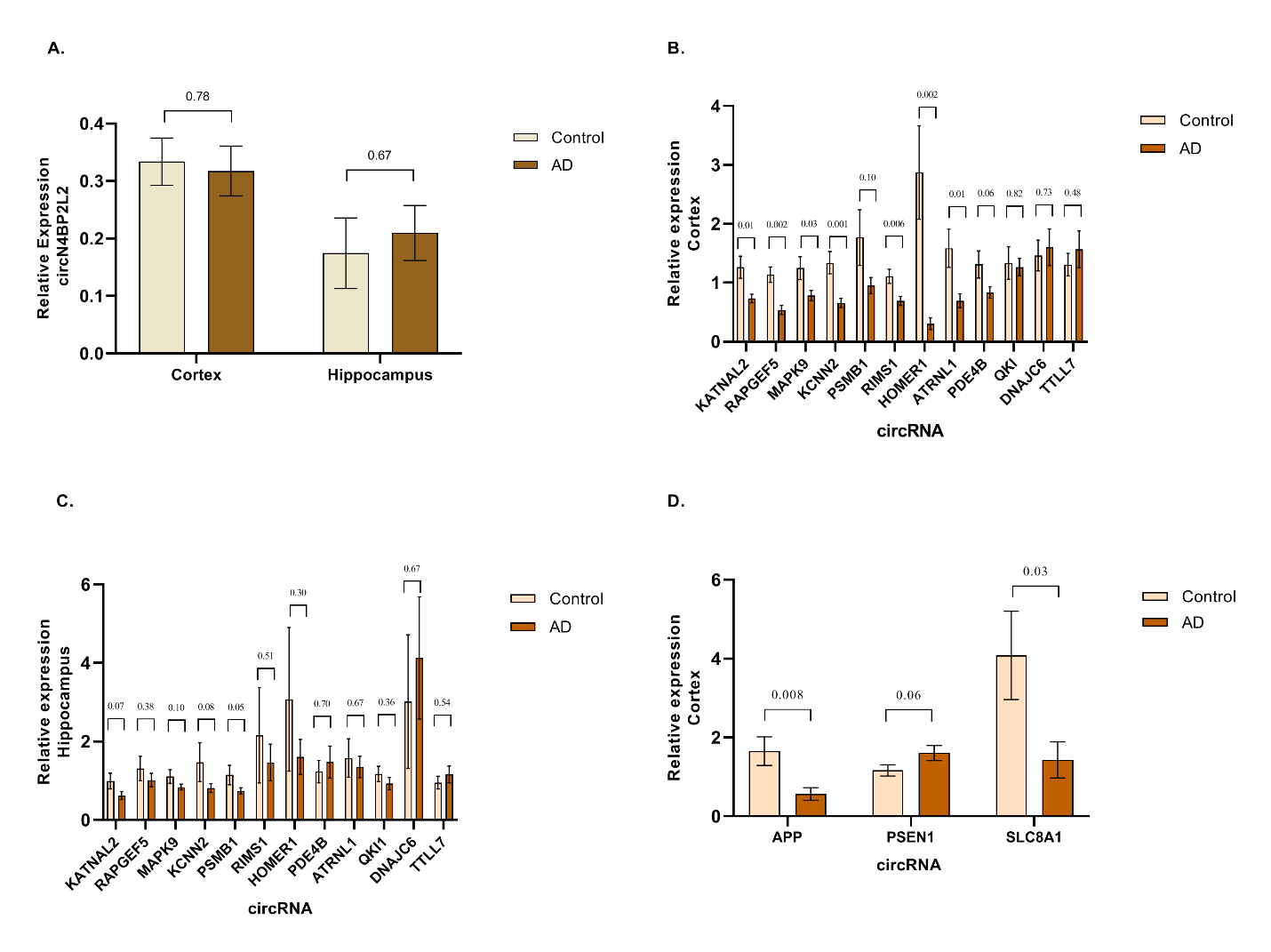

**Supplementary Figure 6**. (A) LDH assay was performed to check for cytotoxicity. oTau treated NPCs exhibit more cell death as compared to vehicle treated. (B) Relative expression of AD associated circRNA does not change when NPCs are subjected to Arsenite stress. Data are given as mean ± SE.

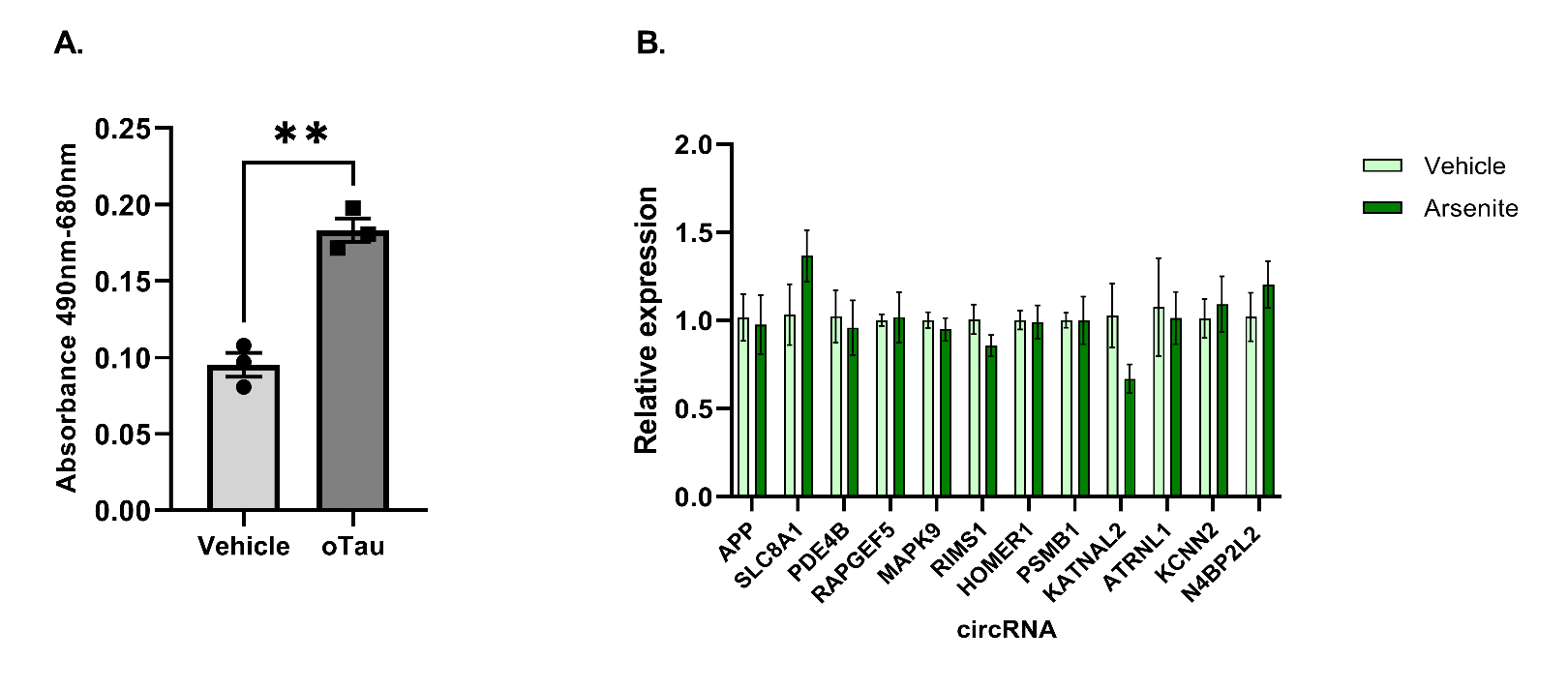
